# Compensatory Evolution to DNA Replication Stress is Robust to Nutrient Availability

**DOI:** 10.1101/2024.10.29.620637

**Authors:** Mariana Natalino, Marco Fumasoni

## Abstract

Evolutionary repair refers to the compensatory evolution that follows perturbations in cellular processes. While evolutionary trajectories are often reproducible, other studies suggest they are shaped by genotype-by-environment (GxE) interactions. Here, we test the predictability of evolutionary repair in response to DNA replication stress—a severe perturbation impairing the conserved mechanisms of DNA synthesis, resulting in genetic instability. We conducted high-throughput experimental evolution on Saccharomyces *cerevisiae* experiencing constitutive replication stress, grown under different glucose availabilities. We found that glucose levels impact the physiology and adaptation rate of replication stress mutants. However, the genetics of adaptation show remarkable robustness across environments. Recurrent mutations collectively recapitulated the fitness of evolved lines and are advantageous across macronutrient availability. We also identified a novel role of the mediator complex of RNA polymerase II in adaptation to replicative stress. Our results highlight the robustness and predictability of evolutionary repair mechanisms to DNA replication stress and provide new insights into the evolutionary aspects of genome stability, with potential implications for understanding cancer development.

## Introduction

Compensatory evolution is a process by which cells mitigate the negative fitness effects of persistent perturbations in cellular processes across generations. This adaptation occurs through the accumulation of spontaneously arising mutations that progressively alleviate cellular defects, ultimately restoring fitness (Kimura, 1985). Perturbations can affect virtually any cellular process and may arise from genetic disruptions due to mitotic errors, the presence of selfish genetic elements, or the cytotoxic effects of external agents such as drugs (LaBar et al., 2020). Consequently, compensatory evolution, often referred to as ‘evolutionary repair,’ is relevant for the evolution of organisms in nature (Natalino & Fumasoni, 2023), as well as in the context of viral (Debray et al., 2022), bacterial (Yang et al., 2020), and parasitic infections (McCutchan et al., 2004), or during cancers’ somatic evolution (Persi et al., 2021). Predicting the evolutionary trajectories that lead to evolutionary repair can shed light on fundamental principles underlying the evolution of cell biology and inform clinical treatments.

Understanding evolutionary outcomes, however, is challenging (Lässig et al., 2017; Papp et al., 2011; Wortel et al., 2021), largely due to the polygenic nature of traits (Fagny & Austerlitz, 2021) and their interaction with the environment (Boyer et al., 2021). Genotype-by-environment (GxE) interactions are well-documented. However, their frequency and impact vary widely among different populations and traits (Fumasoni, 2020; Gorter et al., 2017; Hietpas et al., 2013; Saltz et al., 2018), raising questions about the extent to which the environment shapes evolutionary trajectories during evolutionary repair. A study by Szamecz and colleagues examined the evolutionary trajectories of 180 haploid yeast gene deletions over 400 generations and reported that the accumulation of adaptive mutations in response to gene loss-of-function (LOF) resulted in significant fitness differences across different environments compared to evolved wild-type (WT) controls (Szamecz et al., 2014).

Similarly, Filteau and colleagues found that the carbon source impacted the evolutionary trajectories available to recover from a mutation in the *LAS17* gene, which mimics the causal mutation of Wiskott-Aldrich syndrome in humans (Filteau et al., 2015). On the other hand, a number of studies in yeast have reported a high level of parallelism in evolutionary trajectories, when conserved cellular processes such as chromosome cohesion (Hsieh et al., 2020), segregation (Pavani et al., 2021), and cell polarization (Laan et al., 2015) are perturbed under a stable nutrient-rich environment.

For example, compensatory evolution to constitutive replication stress, an intracellular perturbation that challenges the faithful replication of the genome, was reported to reproducibly depend on adaptive changes in DNA replication, chromosome segregation and cell cycle modules (Fumasoni & Murray, 2020, 2021). Replication stress is considered an early hallmark of cancer (Macheret & Halazonetis, 2015), deriving directly from oncogene activation, and sustaining genetic instability throughout cancer progression (Gaillard et al., 2015). Cancer cells experiencing DNA replication stress can divide in widely variable nutrient environments depending on the site of the primary lesion, its metastatic derivatives, and the level of tumor vascularization (Torrence & Manning, 2018). Several studies in recent years have linked nutrient availability and metabolic processes orchestrated by the Target of Rapamycin (TOR) pathway to genome maintenance processes such as DNA replication (Lamm et al., 2019; Schonbrun et al., 2013; Silvera et al., 2017) and repair (Ferrari et al., 2017; Shen et al., 2007; Shimada et al., 2013). Here we leverage a well-characterized system to induce and monitor adaptation to constitutive DNA replication stress in *Saccharomyces cerevisiae* budding yeast to explore the impact of different physiologically relevant levels of nutrients on evolutionary repair.

We evolved 96 parallel populations of budding yeast under conditions of glucose starvation or abundance, with or without replication stress. Constitutive replication stress was induced by deleting *CTF4*, a non-essential replication fork component responsible for coordinating helicase progression with other fork-associated activities (Gambus et al., 2009; Samora et al., 2016; Villa et al., 2016; Yuan et al., 2019). Glucose availability significantly impacted the growth rate, cell cycle progression, and cell size of replication stress mutants (*ctf4*Δ). Notably, glucose starvation restored some cell cycle traits to WT levels, and improved the competitive fitness of *ctf4*Δ mutants. Unlike WT controls, replication stress mutants evolved faster under higher glucose concentrations. Nevertheless, whole-genome sequencing of evolved *ctf4*Δ populations revealed high genetic robustness and parallelism across glucose conditions. Importantly, we identified a novel module involved in adaptation to replication stress through transcriptional regulation. Overall, our findings demonstrate the predictability and robustness of compensatory evolution to constitutive replication stress and challenge the idea that environmental constraints inevitably lead to distinct evolutionary outcomes. These findings have significant implications for predicting evolutionary trajectories during evolutionary repair, impacting our understanding of the evolution of cellular processes, and of medical conditions such as cancer progression.

## Results

### Glucose availability impacts cell physiology and fitness in the presence of DNA replication stress

The impact of nutrient availability on yeast cell cycle and growth dynamics has been well-documented (Beck & von Meyenburg, 1968; Brauer et al., 2008; Carter & Jagadish, 1978; Johnston et al., 1977, 1980; Slater et al., 1977; Talavera et al., 2024). Reduced nutrient availability typically decreases population growth rates and prolongs the G1 phase, due to the checkpoint controlling entry into the S phase (Alberghina et al., 1998; Johnston et al., 1977; Turner et al., 2012). However, it is still unclear whether these effects are similarly observed in cells experiencing significant intracellular defects, or if they are overshadowed by the cell cycle disruptions these defects cause. In the absence of Ctf4, cells exhibit multiple defects associated with DNA replication stress, such as single-stranded DNA gaps and altered replication forks, leading to cell cycle checkpoint activation and severe growth impairments (Abe et al., 2018; Fumasoni et al., 2015; Miles & Formosa, 1992).

To investigate how glucose availability affects the growth and physiology of *S. cerevisiae* experiencing constitutive replication stress, we grew WT and *ctf4*Δ cells to different glucose abundances ranging from starvation (0.25%) to excess (8%). As the initial glucose concentration decreased, WT cells showed progressively lower growth rates (Fig 1A), longer doubling times (Fig S1A), and reduced maximum optical density (Fig S1B). *ctf4*Δ cells exhibited a similar, albeit smaller, decline (Fig 1A, right), despite their growth rates in standard conditions (2% glucose) being already significantly lower than the minimum observed in starved WT cells. In WT cells, nutrient-dependent reductions in growth rates are linked to changes in cell cycle progression and cell size (Johnston et al., 1979; Turner et al., 2012). We asked how these growth differences affect cell physiology and cell cycle progression in cells experiencing DNA replication stress. Previous studies showed that *ctf4*Δ cells present an altered cell-cycle profile, characterized by a prominent G2 arrest and an almost absent G1 phase (Fumasoni et al., 2015; Tanaka et al., 2009). The extended G2 phase results from the constitutive activation of the DNA damage checkpoint, which delays anaphase to allow for DNA lesion repair (Fumasoni & Murray, 2020; Poli et al., 2012; Tanaka et al., 2009). The shortened G1 phase is usually attributed to the large size of cells arrested in mitosis, leading to premature satisfaction of the size checkpoint during the subsequent G1 phase (Johnston et al., 1977). Similar to WT cells (Fig 1B, left), decreasing glucose availability in *ctf4*Δ cells led to changes in cell-cycle profile (Fig 1B, right), increasing the length of G1 (Fig 1C), reducing that of G2 (Fig S1C), and resulting in smaller cell sizes (Fig 1D). Notably, glucose starvation (0.25%) alleviated these physiological defects in replication stress mutants, leading to a G1 length and a cell size indistinguishable from those of WT cells under standard glucose abundance (6.33 ± 0.7 min vs. 7.40 ± 1.1 min and 4.61 ± 1.5 µm vs. 4.68 ± 0.4 µm, respectively).

**Figure 1.**
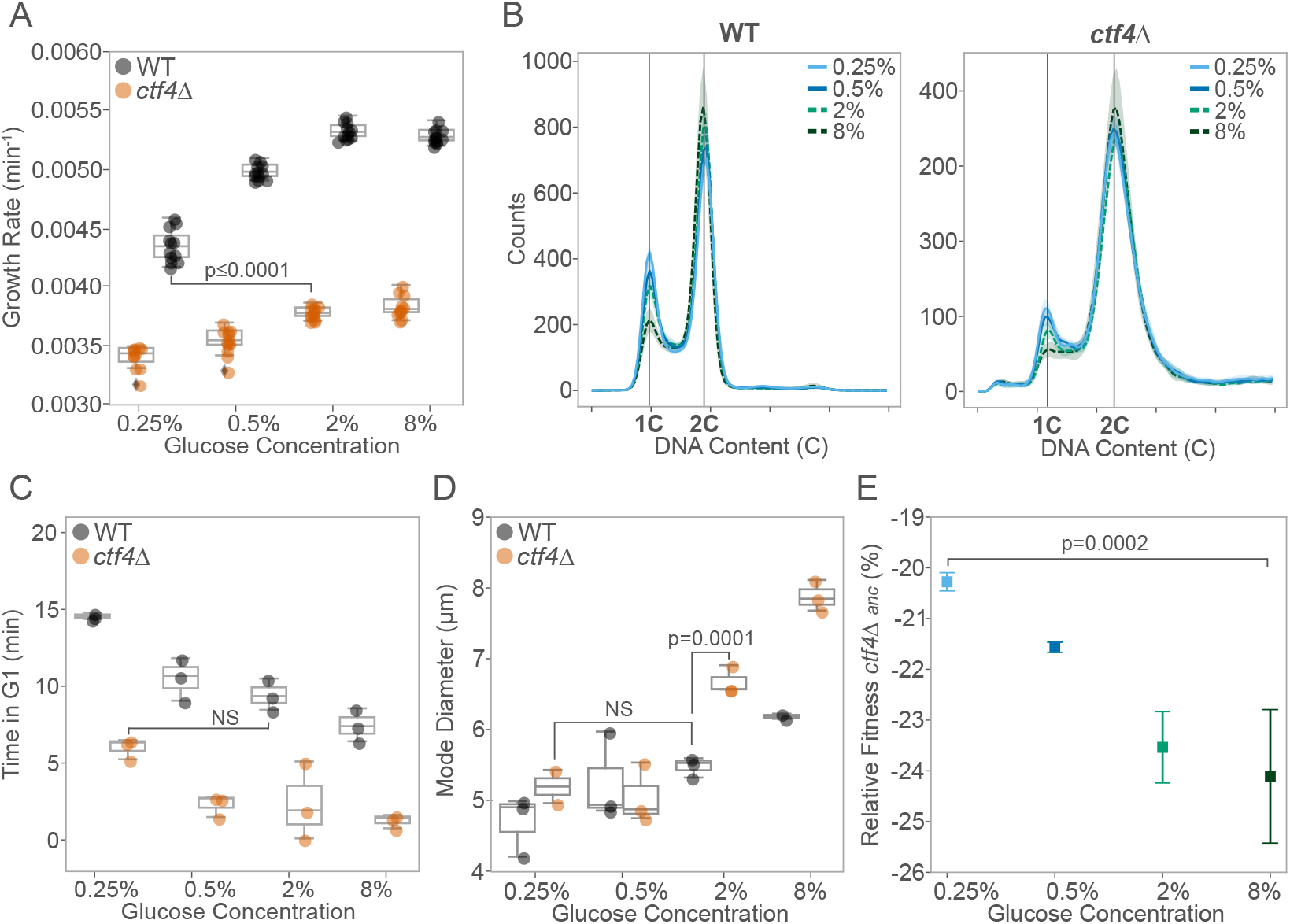
Glucose concentration impacts cell physiology in the presence of DNA replication stress. (A) Population growth rates (min^-1^) of ancestral WT (black) and *ctf4*Δ mutant (orange) across different glucose concentrations. Box plots show median, interquartile range (IQR), and whiskers extending to 1.5×IQR, with individual data points beyond whiskers as outliers. (B) Cell-cycle profiles of ancestral WT (left) and *ctf4*Δ (right) across glucose concentrations. Colors and line styles represent different glucose concentrations: solid line light blue (0.25%), solid line dark blue (0.5%), dash line light green (2%), and dash line dark green (8%). Bold lines indicate mean profiles, and shaded areas represent standard deviation (SD). 1C represents the DNA content of a cell in G1 and 2C of a cell in G2/M. (C) Time spent (minutes) in G1 phase for ancestral WT and *ctf4*Δ, across different glucose concentrations, estimated from DNA content and doubling times (see Materials and Methods). (D) Mode cell diameter of ancestral WT and *ctf4*Δ across different glucose concentrations. (E) Mean relative fitness of ancestral *ctf4*Δ relative to reference WT across different glucose concentrations. Colors represent glucose concentration. Error bars represent SD. Detailed statistical analysis and underlying data for this figure are provided in Supplementary File 1.

We assessed the impact of these cell cycle changes on cellular fitness using competition assays, in which two genetically distinct lineages compete for nutrients in the same media over multiple generations (see Materials and Methods for details). As previously reported, *ctf4*Δ cells exhibited severe fitness defects (Fumasoni & Murray, 2020, 2021). However, their relative fitness improved compared to the WT reference as the initial glucose levels in the competition media decreased. In 0.25% glucose, *ctf4Δ* cells showed approximately a 4% fitness advantage over those competed in 8% glucose (Figure 1E). These results highlight how changes in glucose availability impact the population growth and cell physiology of *ctf4*Δ cells, mitigating the fitness defects introduced by constitutive DNA replication stress.

### Replication stress mutants display higher adaptation rates and fitness gains under glucose abundance

Can glucose availability influence the evolutionary adaptation to DNA replication stress? The glucose-dependent effects on fitness and cell physiology reported above alter the selective pressures experienced by populations. Over multiple generations, these could influence the cells’ ability to evolutionarily repair DNA replication stress. To test this hypothesis, we evolved 12 parallel populations each of haploid *ctf4*Δ mutants and WT controls over 1000 generations through daily serial dilutions in rich media (YP) under glucose starvation, or glucose abundance (Fig 2A). Importantly, we adjusted the number of cells passaged and the daily number of generations to maintain the effective population size (Ne) at the same order of magnitude throughout the experiment (Fig S2A). Maintaining a stable Ne is crucial, as large differences can influence evolutionary dynamics by affecting genetic drift and the population’s access to beneficial mutations, potentially confounding the impact of nutrient levels in adaptation (Crow & Kimura, 1970; Desai et al., 2007; Lynch et al., 1995; Schenk et al., 2022; Silander et al., 2007; Van den Bergh et al., 2018; Wright, 1931).

**Figure 2.**
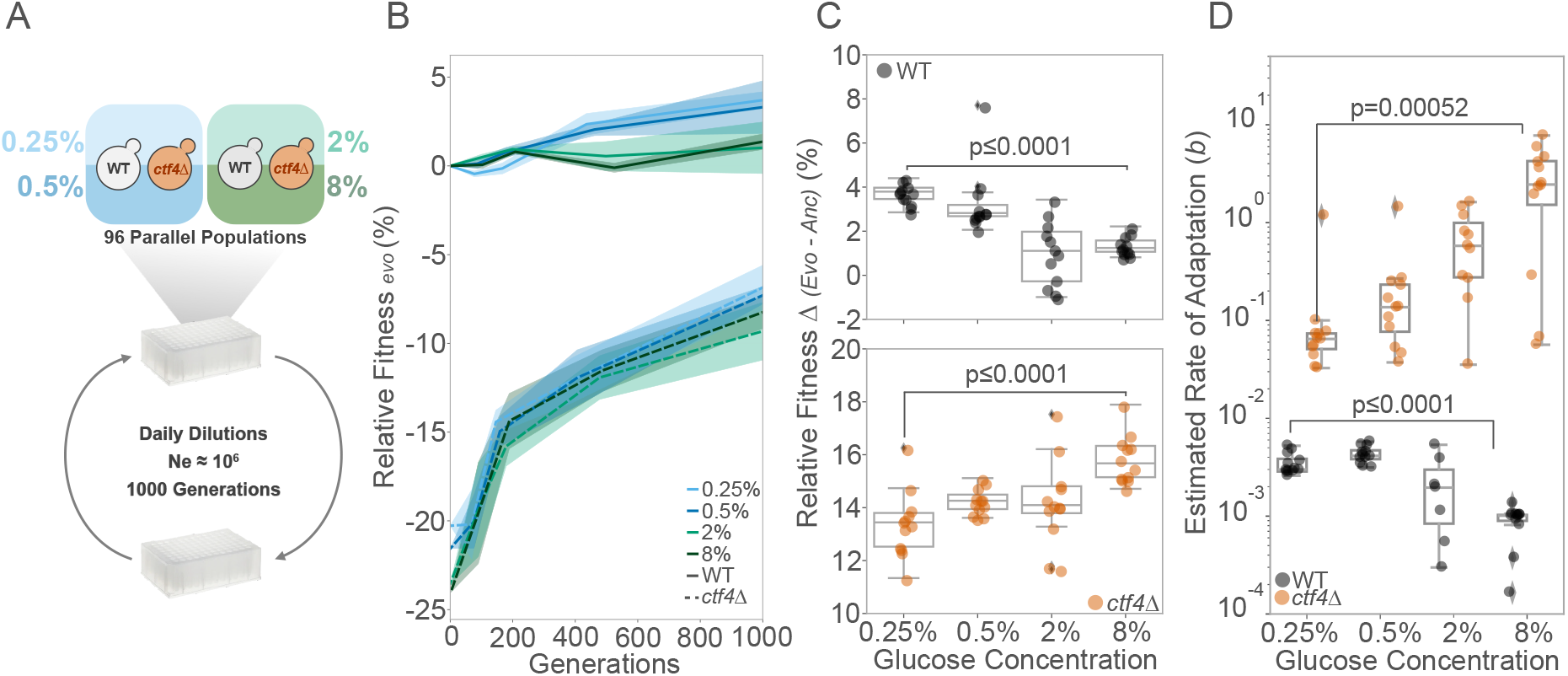
Glucose availability impacts the dynamics of fitness recovery. (A) Schematic of experimental layout. 48 isogenic clones of ancestral *ctf4*Δ and 48 WT clones were inoculated in either glucose starvation (0.25% and 0.5%) or abundance (2% and 8%) on deep 96-well plates. Clones were grown to saturation, and diluted daily until reaching 1000 generations. Bottleneck was adjusted to maintain Ne within the same order of magnitude throughout the experiment. (B) Relative fitness trajectories relative to a WT reference over 1000 generations. Solid and dashed lines indicate mean fitness of WT and *ctf4*Δ evolved populations (1000 generations), respectively. Shaded areas represent the standard error of the mean (SEM) across the 12 parallel populations. Colors represent glucose concentrations: light blue (0.25%), dark blue (0.5%), light green (2%), and dark green (8%). (C) Fitness gains (Δ) at generation 1000 for evolved WT (upper panel, black) and *ctf4Δ* (lower panel, orange) populations. Box plots show median, IQR, and whiskers extending to 1.5×IQR, with individual data points beyond whiskers considered outliers. Fitness gains were calculated by subtracting ancestral relative fitness from evolved populations’ relative fitness, per glucose concentration. (D) Estimated rate of adaptation (parameter *b*) across glucose concentrations. Each datapoint represents the estimated rate of adaptation of a single evolved population of either WT or *ctf4*Δ. Detailed statistical analysis and underlying data for this figure are provided in Supplementary File 2.

By generation 1000, both WT and *ctf4*Δ evolved lines achieved, on average, slightly higher fitness in low glucose compared to high glucose conditions (Fig 2B, FigS2B). However, due to the varying initial fitness of *ctf4*Δ cells across different glucose environments, they display an opposite trend to WT, with increasing absolute fitness over the course of the experiment as glucose concentration rose (Fig 2C). We further looked at the fitness dynamics of the evolved lines across the experiment (Fig 2B). All fitness trajectories were approximated by a power law function, which has been proposed to model long-term evolutionary dynamics in constant environments (Wiser et al., 2013). The parameter *b* of the power law function represents the rate of adaptation, with higher values of *b* indicating a faster adaptation rate. We used this parameter to quantify and compare the adaptation rates of each lineage throughout the evolution experiment. The adaptation rates in WT and *ctf4*Δ cells showed opposing trends in response to glucose availability. WT cells adapted faster in low glucose, while *ctf4*Δ lines showed progressively higher adaptation rates as glucose levels increased (Fig 2D). These results demonstrate that cells can recover from fitness defects caused by constitutive DNA replication stress regardless of the glucose environment. However, we show that adaptation rates under DNA replication stress exhibit opposing trends compared to WT cells, with faster adaptation resulting in greater fitness gains in higher glucose conditions.

### Genotype, and not the environment, shapes the mutational profile in evolved populations

During experimental evolution, populations accumulate various types of mutations: adaptive mutations that confer a fitness benefit and neutral or slightly deleterious mutations that spread due to genetic drift or hitchhiking alongside adaptive mutations. Comparative analysis of the identity and frequency of these mutations can shed light on the genetic basis of the evolutionary processes across different environments. We whole-genome sequenced the final evolved populations as well as the ancestral clones (see Materials and Methods), to examine whether genotype, environment, or their interaction (GxE) affected the mutational landscape of the evolved lines. Evolved *ctf4*Δ lines exhibited approximately 3 to 4 times more mutations in coding regions (CDS) than WT lines (Fig 3A). This finding suggests that replication stress mutants either accumulate mutations at a higher frequency than WT or that a larger fraction of mutations were beneficial and thus were retained until generation 1000. To distinguish between these scenarios, we examined the frequency of synonymous mutations, which are less likely to be subject to selection during evolution. We found no significant differences in synonymous mutations between WT and *ctf4*Δ populations (Fig S3A). These results support the hypothesis that replication stress in *ctf4*Δ lines favors the retention of beneficial mutations, rather than simply increasing the overall mutation rate. We detected a higher median mutation fraction (Fig S3B) and significantly different distribution (p ≤ 0.0001) in *ctf4*Δ compared to WT evolved populations, with a strongly skew towards higher-frequency mutations (Fig 3B). This indicates a higher level of clonality in *ctf4*Δ populations, consistent with a greater frequency of adaptive mutations reaching fixation. Despite significantly affecting cell physiology and fitness dynamics, glucose concentrations did not influence mutation counts (Fig 3C) or population clonality (Fig S3C) in either WT or *ctf4*Δ evolved lines.

**Figure 3.**
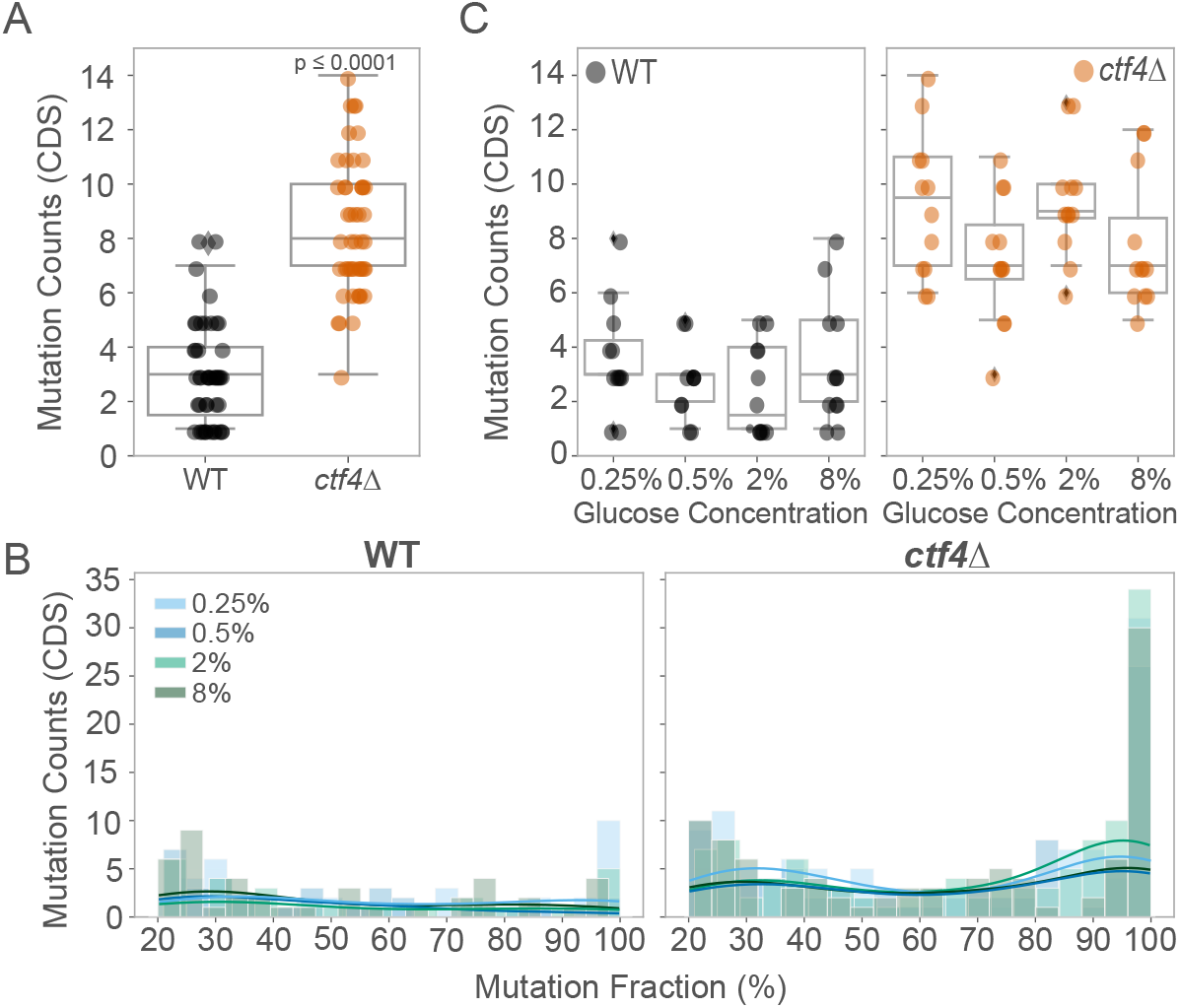
The mutational profile is mainly influenced by the genotype. (A) Total detected mutations in CDS per evolved WT (black) and *ctf4*Δ (orange) populations, at generation 1000. Box plots show median, IQR, and whiskers extending to 1.5×IQR, with individual data points beyond whiskers considered outliers. (B) Distribution of mutated read fractions across glucose concentrations for WT (left) and *ctf4Δ* (right) evolved populations, used as a proxy for clonality. Mutation fraction (%) was calculated as the fraction of reads from whole populations sequencing that contained a particular mutation in CDS. Colors represent glucose concentrations: light blue (0.25%), dark blue (0.5%), light green (2%), dark green (8%). Histograms are overlaid with a kernel density estimate (KDE, colored lines) to illustrate frequency distributions. Kolmogorov-Smirnov (KS) tests were used to compare read fraction distributions between glucose concentrations and genotypes. (C) Total mutations detected in CDS across glucose concentrations for WT (left) and *ctf4Δ* (right) populations at generation 1000. Detailed statistical analysis and underlying data for this figure are provided in Supplementary File 3.

We then asked whether glucose concentrations influenced the occurrence of putative adaptive mutations within each genotype. We identified candidate adaptive mutations by determining which genes were mutated more frequently than expected by chance in independent populations (see Materials and Methods for details). In WT, 13 mutations across 48 evolved populations exhibited signs of selection, with 7 of these mutations occurring exclusively under glucose starvation (Fig 4A, upper panel). Only *MSS11*, encoding for a transcription factor involved in regulating filamentous growth, was found to be mutated across all glucose conditions (Fig 4A, upper panel). All positively selected GO terms, including genes involved in nutrient sensing and cell growth—such as those in the Mitogen-Activated Protein Kinase (MAPK) signaling pathway and the STErility gene family—were exclusively found in populations evolved under low glucose, but not in other glucose concentrations (Fig 4A, bottom panel). In *ctf4*Δ populations, we similarly identified more putatively adaptive genes uniquely mutated under glucose starvation (0.25% and 0.5%) compared to abundance (2% and 8%) (11/20 vs 4/20, Fig 4B, upper panel). However, GO term enrichment analysis revealed a different pattern compared to WT lines. While only a few modules involved in rRNA regulation and MAPK signaling were exclusively selected under glucose starvation, most selected GO terms – such as those involved in cell cycle, transcription regulation and genome maintenance – were found enriched across all glucose conditions (Fig 4B, bottom panel, and 4C).

**Figure 4.**
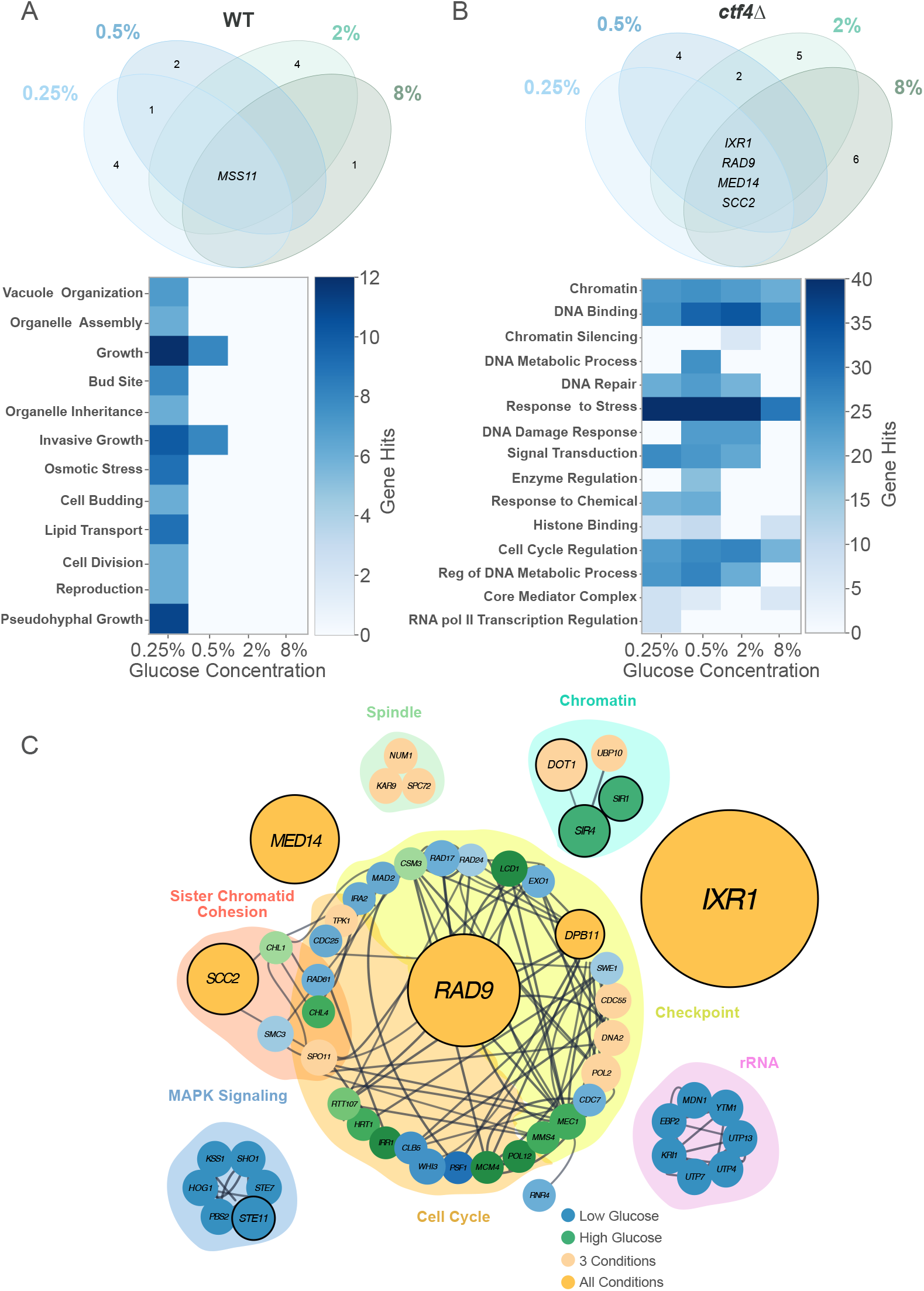
Environment impacts the genetic basis of adaptation of WT but not replication stress mutant. (A) Venn diagram of putative adaptive genes mutated in evolved WT populations across glucose concentrations (upper panel). Colors represent glucose concentrations: light blue (0.25%), dark blue (0.5%), light green (2%), and dark green (8%). Numbers represent counts of putative adaptive genes (excluding zero counts). Gene names in the center are shared across conditions. GO-term enrichment across glucose concentrations in WT (bottom panel). Heatmap illustrates the total number of gene hits for a significant GO-terms. Fisher’s Exact Test was used to assess significance between pairwise glucose conditions. (B) Venn diagram of putative adaptive genes mutated in evolved *ctf4*Δ populations across glucose concentrations (upper panel) and corresponding GO-term enrichment heatmap (bottom panel). (C) Simplified interaction network of mutations detected in *ctf4*Δ evolved populations. Dark grey lines represent known genetic and physical interactions from the literature (https://string-db.org). Node diameter is proportional to the number of populations with mutations in that gene. Nodes are color-coded: blue for mutations in low glucose (0.25% and 0.5%), green for high glucose (2% and 8%), light orange for both high and low glucose, and dark orange for all conditions. Nodes with a bold outline indicate putative adaptive genes. Shaded clusters represent GO term enrichment for biological processes obtained using STRING. Network was curated in Cytoscape. Detailed statistical analysis and underlying data for this figure are provided in Supplementary File 3.

In particular, four genes, here defined ‘core genetic adaptation’ were consistently mutated (three SNPs/INDELS and one gene amplification) in response to replication stress regardless of glucose availability (Fig 4B and Supplementary File 3). Three of these genes had been previously identified to affect DNA replication (*IXR1*), the DNA damage checkpoint (*RAD9*), and chromosome cohesion (*SCC2*) in response to constitutive DNA replication stress (Fumasoni and Murray 2020 and 2021). Importantly, we identified a novel fourth module, represented by mutations in *MED14*, also known as *RGR1*, a gene implicated in transcriptional regulation (Sakai et al., 1988; Warfield et al., 2022). Overall, our results indicate that the mutational profile of replication stress mutants is primarily shaped by the intracellular stress imposed by the *CTF4* deletion. The genotype, rather than the glucose environment, played a dominant role in determining the mutational profiles. Additionally, while glucose availability influenced adaptive mutations in WT, this effect was negligible under DNA replication stress, where most selected modules were consistently targeted across all conditions.

### A Single-Amino Acid Substitution in Mediator Complex Subunit 14 Alleviates Replication Stress Defects

We observed a remarkable level of parallelism targeting the Med14 subunit of the transcription mediator complex in *ctf4*Δ evolved populations. A single amino acid substitution of histidine with proline in the C-terminal of the protein (*med14*-H919P) was detected at high frequency in 22 out of 48 of independently evolved populations (Fig 5A). The reconstruction of this mutation in the *ctf4*Δ ancestor demonstrated causality by leading to approximately a 4% fitness advantage (Fig 5B).

**Figure 5.**
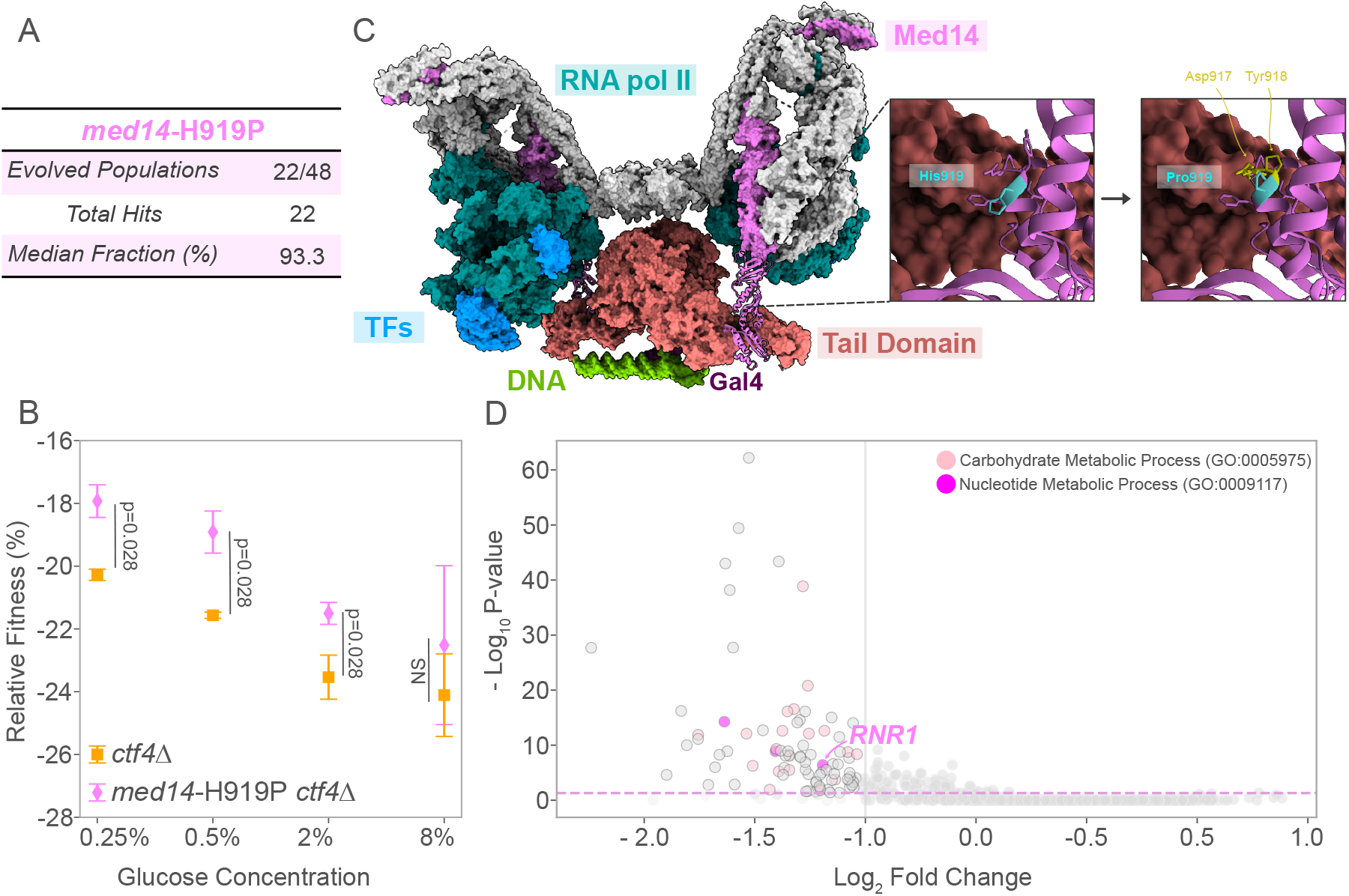
Adaptive fitness and structural insights of Med14 mutation in replication stress mutants. (A) Schematic representation of the prevalence of the *med14*-H919P point mutation in evolved *ctf4Δ* populations across glucose conditions. (B) Mean relative fitness of *ctf4*Δ ancestor (orange) and *ctf4*Δ carrying reconstructed mutation *med14*-H919P (pink). Error bars represent SD. (C) Composite model of the transcription pre-initiation complex of RNA pol II with mediator complex forming a dimer to act on a distal promoter (Gal4-activated) (PDB: 7UIO). Tail components, RNA pol II, transcription factors (TFs), DNA, and regulatory Gal4 are color-coded. Med14 is highlighted in pink, with secondary structures shown for the C-terminal domain (705-1082 aa). Right panel: top view of HIS 919 (blue) and surrounding amino acids, with a simulation of the HIS to Pro substitution at site 919 using Chimera X. Structural clashes were identified using Chimera X (affected residues in yellow). (D) Volcano plot of transcriptional changes after degron-removal of Med14 C-terminal (Warfield et al., 2022). Purple dashed lines indicate the significance threshold (p-value ≤ 0.01). Differentially expressed genes were defined by an adjusted p-value < 0.01 and an absolute log2 fold change > 1 (equivalent to a 2-fold change). GO enrichment analysis of downregulated genes shows enrichment in carbohydrate (light pink) and nucleotide (magenta) metabolic processes. Detailed statistical analysis and underlying data for this figure are provided in Supplementary File 4 and 5.

The mediator complex is essential for the initiation of eukaryotic gene transcription (Myers et al., 1998; Thompson & Young, 1995) and is highly conserved across eukaryotes (Bourbon, 2008). It interacts with the C-terminal domain of RNA polymerase II (RNA pol II) and transcription factors, facilitating transcription initiation (Kim et al., 1994). In yeast, the mediator complex comprises 25 subunits, organized into three distinct modules: kinase, core mediator, and the tail. The tail module plays a role in gene-specific transcription regulation by interacting with transcription activators (Soutourina, 2018). Recent structural studies depicted the dimerization of yeast’s pre-initiation complex upon binding to divergent promoters (Fig 5, left; PDB: 7UIO) (Gorbea Colón et al., 2023). The C-terminal domain of Med14, known as Tail Interaction Domain (TID, residues 705-1,082), links the tail module to the remaining mediator complex. The adaptive mutation *med14*-H919P is located within the TID, specifically in an alpha-helix that connects Med14 with the core mediator complex (Fig 5C, left). We simulated Med14’s amino acid substitution using the Rotamer tool in Chimera X, which predicted steric clashes with adjacent amino acids, Asp 917 and Tyr 918 (Fig 5C, right). Additionally, the substitution of histidine with proline is predicted to destabilize the alpha-helix due to proline’s inability to form hydrogen bonds with neighboring residues, suggesting a possible disruption of Med14’s structural integrity. To explore possible functional consequences of this mutation, we analyzed the publicly available RNA-seq data resulting from degron-mediated removal of the Med14 TID (Warfield et al., 2022). Our analysis identified 90 significantly downregulated genes, with an enrichment in energy and nucleotide metabolism pathways (Supplementary File 4). The list included a subunit of the ribonucleotide reductase (Rnr1), the enzyme responsible for producing the deoxyribonucleotides (dNTPs) required for DNA replication (Fig 5D). Overall, these results show how an amino acid substitution in the Med14 subunit of the mediator complex, putatively affecting transcription, is strongly selected, and advantageous, in the presence of constitutive DNA replication stress.

### Core genetic adaptation to replication stress is robust to nutrient availability

The signs of positive selection towards the four genes belonging to the core genetic adaptation do not necessarily imply that their fitness benefit does not depend on the glucose environment. To test this point, we engineered loss-of-function alleles mimicking the mutations affecting *IXR1* and *RAD9*, as well as the *SCC2* amplification and *med14*-H919P substitution, into the ancestral WT and *ctf4*Δ cells. We competed all reconstructed strains individually against a reference WT strain and assessed their relative fitness under the different glucose environments. In the WT background, all mutations mostly resulted in neutral or slightly deleterious effects of fitness (Fig S4A). When introduced in a *ctf4*Δ background, all mutations led to fitness benefits compared to the ancestral cells. However, with the exception of *IXR1* deletion, competition assays performed in the different glucose conditions resulted in comparable fitness benefits with no statistical difference between glucose starvation or abundance (Fig S4B). Similarly, we did not detect differences in the frequency of occurrence or average fractions achieved by the mutations in the populations evolved under different glucose environments (Fig S4C). The presence of all mutations in the final evolved lines correlated with their fitness benefits, suggesting how their selection in all glucose conditions was mostly dictated by their relative fitness benefits (Fig 6A). Is the combined effect of the mutations belonging to this core genetic adaptation sufficient to recapitulate the final fitness of the evolved *ctf4*Δ populations? (Fig S2B). To address this question, we computed the cumulative fitness benefit, assuming additivity, of the core adaptation mutation present in each evolved population (Supplementary File 5). For all glucose concentrations, the expected fitness closely approximated or exceeded the final fitness of the evolved populations (Fig 6B). Overall, these results demonstrate the robustness of the core genetic adaptation to DNA replication stress, represented by four mutations, recurrently selected in all conditions based on their fitness benefits. Together, these mutations largely explain the fitness gains observed in populations evolved over 1000 generations. We then investigated the extent to which the core genetic adaptation to DNA replication stress was beneficial under alternative nutrient conditions. We reconstructed a quadruple mutant in the ancestral *ctf4*Δ background carrying a frequent combination of adaptive mutations (*ixr1*Δ, and *rad9*Δ and 2x*SCC2*). We then compared its relative fitness across conditions of starvation or abundance of the essential macronutrients’ glucose, nitrogen, and phosphate. Across all conditions, the reconstructed strain showed a significant fitness recovery compared with ancestor *ctf4*Δ (Fig 6C). Altogether, these results demonstrate how the core genetic adaptation we identified is responsible for a large extent of fitness benefits across a wide range of macronutrient availability and, thus, largely robust to nutrient environments.

**Figure 6.**
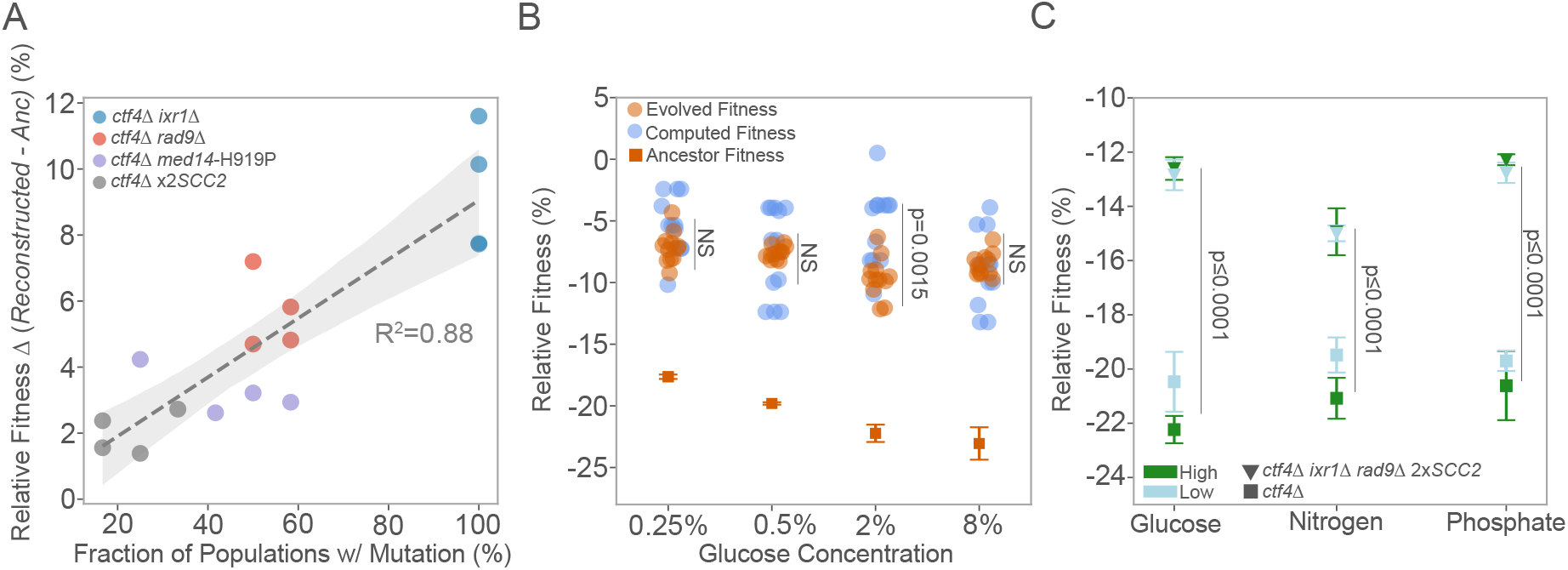
Robustness in core genetics of adaptation to replication stress. (A) Correlation between the fraction of populations carrying specific adaptive mutations and associated fitness benefits (Δ) under replication stress. Each point represents a mutation, color-coded by gene: *ixr1*Δ (blue), *rad9*Δ (red), *med14*-H919P (purple), and 2x*SCC2* (gray). The x-axis shows the fraction of evolved populations with each mutation, while the y-axis shows the conferred fitness advantage in the ancestral *ctf4Δ* background (%) for each glucose concentration. A positive correlation (R^2^ = 0.88) is observed, with the shaded area representing the 95% confidence interval of the linear regression. (B) Comparison of ancestral, evolved, and computed fitness for *ctf4Δ* lines. For each evolved population, if a reconstructed gene was found mutated, its fitness effect in the respective glucose concentration was added to the *ctf4Δ* ancestor to calculate computed fitness (blue). Relative fitness of evolved *ctf4Δ* populations (orange) and ancestral *ctf4Δ* (dark orange) are shown. Error bars represent SD. (C) Mean relative fitness of *ctf4*Δ and reconstructed strain (*ctf4*Δ *ixr1*Δ *rad9*Δ 2x*SCC2*). Error bars represent SD. Each point indicates the mean fitness for a given nutrient and concentration (low or high), with marker shapes differentiating the genotypes (squares for *ctf4*Δ and inverted triangle for reconstructed) and colors representing concentration levels (light blue for low and dark green for high). Detailed statistical analysis and underlying data for this figure are provided in Supplementary File 5.

## Discussion

Gaining a deeper understanding of gene-by-environment (GxE) interactions within the context of compensatory evolution is crucial for elucidating how cells adapt and maintain functionality in the face of genetic and environmental pressures. In this work, we investigated how changes in glucose availability, a common environmental variable, shaped the evolutionary repair of DNA replication stress. We examined how glucose impacted (i) key cellular phenotypes (ii) the dynamics of fitness recovery, (iii) the mutational profile and (iv) the genetics of adaptation in the presence or absence of DNA replication stress.

### Glucose availability affects cell physiology, fitness and evolutionary dynamics

Glucose availability significantly impacts the growth dynamics of both WT and *ctf4*Δ cells. In our study, glucose starvation partially restored cell cycle and the cell size of *ctf4*Δ cells to levels similar to WT, potentially explaining the proportional decline in fitness of *ctf4*Δ ancestor as glucose levels increased. Several explanations could account for this observation. One possibility could be the reduction in the nucleotides (dNTP) pool under glucose starvation. The reduced activity of the ribonucleotide synthetase, the enzyme responsible for dNTP production has been shown to slow down replication fork progression (Poli et al., 2012) and improve the overall fitness of *ctf4*Δ cells (Fumasoni & Murray, 2020; Poli et al., 2012). During glucose starvation the levels of Rnr1, a subunit of the ribonucleotide reductase, decrease substantially (Corcoles-Saez et al., 2019), likely causing a subsequent drop in dNTP levels. Additionally, *IXR1* loss-of-function, which is strongly selected in evolved *ctf4*Δ lines, results in reduced dNTP pools through reduced expression of *RNR1* (Tsaponina et al., 2011). Interestingly, we observed a significant reduction in the fitness benefits of *IXR1* deletion under glucose starvation (Fig. S4B), suggesting that the fitness effects of *ixr1*Δ may overlap with those of glucose starvation. Another possible explanation involves the alleviation of ribosomal DNA (rDNA)-related replication stress. The rDNA locus is particularly susceptible to DNA replication stress (Salim et al., 2017), with replication fork stalling and collapsing observed, even in WT conditions (Takeuchi et al., 2003). Glucose starvation dramatically reduces rDNA origin firing by 80% in extreme glucose starvation (0.05%) (Kwan et al., 2013). We speculate that the fitness recovery observed in *ctf4*Δ mutants under starvation could result from a combination of a reduced dNTP pool and reduced fork collapse at the rDNA locus.

Beyond the immediate physiological effects, glucose availability also influenced the evolutionary dynamics of replication stress mutants. Specifically, we found that *ctf4*Δ lines evolved more rapidly during the early generations under glucose-rich conditions. However, despite this accelerated gain in fitness, there were no significant glucose-dependent differences in their mutational profiles. This suggests that initial fitness levels, rather than glucose availability itself, may have driven the differences in adaptation rates (Couce & Tenaillon, 2015).

### A novel mechanism of adaptation to DNA replication stress through the transcription mediator complex

A single amino acid substitution at position 919 of Med14 – an essential component of the mediator complex – was selected in 22 out of the 48 *ctf4*Δ populations across glucose conditions. This mutation became fixed in most populations by generation 1000, indicating strong positive selection (Fig S4C, 5A). The recurrence of this mutation and the fitness advantage conferred suggest a significant role for *med14*-H919P in coping with replication stress. Further analysis predicted that this amino acid substitution could lead to perturbations in the structure of Med14 TID. Analysis of the transcriptome dataset available for the degron-mediated depletion of the Med14 TID revealed the downregulation of *RNR1*, among other genes implicated in energy and nucleotide metabolism (Fig 5D). When introduced in a WT background, *med14*-H919P displayed fitness costs specifically under glucose abundance (Fig S4A). These observations are compatible with reduced dNTP availability in *med14*-H919P cells due to the downregulation of *RNR1*. This condition will, in turn, be detrimental when proliferation rates are high (as in WT in high glucose) but beneficial under constitutive DNA replication stress (*ctf4*Δ). Future work should focus on the transcriptomic analysis and quantification of dNTP levels of *med14*-H919 mutants alone and in combination with DNA replication stress. Importantly, dysregulation of the mediator complex has been implicated in multiple cancer types, and human Med14 has been specifically found to be downregulated in lymphoma (Syring et al., 2016).

### Reproducibility of evolutionary repair

Is compensatory evolution shaped by the environment where cells grow and divide? Our study suggests that adaptive strategies to cope with the severe intracellular cellular stress caused by replication perturbations are both predictable and largely independent of nutrient availability. This finding is particularly surprising considering the several reported links between starvation, the TOR pathway, and genome stability (He et al., 2021; Weisman et al., 2014). Additionally, the effects of glucose on the cell physiology and fitness of replication stress mutants reported here could have suggested different selective pressures during evolution. What could explain the discrepancies between our results, and previous studies highlighting the role of the environment in shaping evolutionary trajectories in the context of evolutionary repair (Filteau et al., 2015; Szamecz et al., 2014)? We propose several non-mutually exclusive hypotheses. First, environmental variability can influence population size, affecting access to and the spread of beneficial mutations. In our experiment, we maintained a consistent effective population size across conditions to control for this variable. Second, qualitative changes in the environment, rarely encountered in nature, may impose more severe constraints on evolutionary trajectories than the quantitative and physiological changes we explored. Third, the environment’s influence on compensatory evolution may depend on the specific cellular module perturbed and its genetic interaction with other modules that are significantly affected by environmental conditions. The actin cytoskeleton, which must rapidly adapt to extracellular stimuli, could be increasingly subjected to interactions with the environment when compared to DNA replication machinery, which is confined to the nucleus. Finally, the severity of the fitness defects caused by cellular perturbations may dictate the robustness of evolutionary adaptation. The stronger the initial perturbation, the more likely it is that adaptive mutations accessible through the genetic interaction network are limited in number and have large effects, making it less likely that the network is significantly influenced by environmental changes. A number of recent findings are relevant to these hypotheses: a recent study proposed how the global yeast genetic interaction network is largely robust to environmental perturbation (Costanzo et al., 2021). An increasing body of work has also proposed how the effect of any given mutation is largely dependent on the fitness of the strain in which it occurs (Couce & Tenaillon, 2015; Johnson et al., 2023). This phenomenon has been explained by invoking global epistasis, defined as the high-order interaction network of the mutation with multiple genetic loci (Diaz-Colunga et al., 2023; Kryazhimskiy et al., 2014; Reddy & Desai, 2021), which has been recently shown to be robust to the environment (Ardell et al., 2024). Future experiments, however, will be needed to discriminate between the above-mentioned hypotheses in the context of evolutionary repair.

Our findings demonstrate that while glucose availability significantly affects the physiology and adaptation speed of cells under replication stress, it does not alter the fundamental genetic mutations that drive fitness recovery and evolutionary repair. The consistency of genetic adaptations across different glucose conditions, and their fitness benefits across different environments, suggest that cells follow a conserved pathway to recover from replication stress. These insights contribute to our broader understanding of the fundamental principles of evolutionary cell biology and hold significant implications for predicting evolutionary outcomes in critical areas such as cancer progression and antibiotic resistance. A deeper understanding of the robustness and predictability of adaptive responses could guide the development of therapeutic strategies targeting environment-independent mechanisms, making them more broadly effective.

## Material and Methods

### Strains

All strains were derivatives of a modified version (Rad5+) of *S. cerevisiae* strain W303 (leu2-3;112 trp1-1; can1-100; ura3-1; ade2-1; his3-11,15; RAD5+) kindly provided by Dana Branzei. All strains and respective genotypes used in this study are listed in Supplementary File 6. The ancestors of WT and *ctf4*Δ strains were obtained by sporulating a *CTF4*/*ctf4*Δ heterozygous diploid. This was done to minimize the selection acting on the ancestor strains before the beginning of the experiment. Diploid strains were grown on YP (1% Yeast-Extract, 2% Peptone) + 2% D-Glucose (YPD), transferred to 2 mL sporulation media (YP + Potassium Acetate (KAc) 2% enriched with adenine and tryptophan (A+T)) and grown for over-night (O/N) at 30°C. Cells were then washed twice in sterile milli-Q water, resuspended in 2% Kac in sterile milli-Q with A+T, and incubated for four days at 25°C. Tetrads were re-suspended in water containing zymolyase (Zymo Research, Irvine, CA, US, 0.025 µ/µL), incubated at 37°C for 1m 45s, and dissected on a YPD plate using a micro-manipulator. Spores were allowed to grow into visible colonies and genotyped by the presence of genetic markers and PCR.

### Media and Growth Conditions

All experiments were conducted in standard rich medium (YP) unless otherwise noted. For glucose availability experiments, D-glucose concentrations were adjusted to 0.25%, 0.5%, 2%, or 8%, using a 50% autoclaved sterile Milli-Q water D-glucose stock solution. YP was supplemented to a final concentration of 0.01% of Adenine and Tryptophan. Transformation experiments were carried out in YP medium supplemented with 2% D-glucose. All cultures were incubated at 30°C. For macronutrient availability experiments, synthetic complete media was used. The composition of synthetic media was as follows: 0.67% Yeast Nitrogen Base (YNB), supplemented with 0.025 mg/mL each of histidine, uracil, tryptophan, and adenine; 0.05 mg/mL leucine; 0.768 mg/L of arginine, isoleucine, lysine, phenylalanine, and tyrosine; 1.152 mg/L aspartic acid, 0.256 mg/L methionine, 1.536 mg/L threonine, 2.176 mg/L valine, and 2% D-glucose. For glucose-limitation, glucose was added to a final concentration of 0.25%. Phosphate and nitrogen availability were adjusted following the conditions described by (Boer et al., 2008). Phosphate and nitrogen restriction and abundance media was prepared with 0.17% YNB without potassium phosphate monobasic and 0.17% YNB without ammonium sulfate, respectively.

### Growth dynamics

Ancestral clones of WT and *ctf4Δ* strains were grown to saturation in YPD (2% Glucose) and diluted 1:100 into 150 µL of YP medium containing different glucose concentrations. Six replicates per condition in a 96-well plate. The plate was covered with breathable membranes (Breath-Easy®, Sigma-Aldrich) and incubated at 30°C for 48 hours in a microplate reader (BioTek Epoch 2, Agilent). Optical density at 600 nm (OD600) was measured every 10 minutes after 10 seconds of orbital shaking (speed). Protocol was defined using Agilent Gene5 software (version 3.12). Growth data were analyzed in Python v3.9 and a script based on growth_curve_analysis.py (https://github.com/nwespe/OD_growth_finder) was adapted. Statistical comparisons between genotypes and glucose conditions were performed using a two-way analysis of variance (ANOVA) with Turkey Honestly Significant Difference (HSD) post-hoc test for pair-wise comparisons. Statistical significance was determined at adjusted p-value ≤ 0.05.

### Cell cycle Analysis

Cell cycle analysis was performed as described by (Fumasoni et al., 2015). Briefly, three independent cultures per genotype of exponentially growing cells (∼1×10^7^ cells) were collected from each glucose condition (0.25%, 0.5%, 2%, or 8%) by centrifugation and incubated in 70% ethanol, then eluted in 250 mM Tris-HCl (pH 7.5) at 4°C overnight. Cells were washed with 50 mM Tris-HCl (pH 7.5), resuspended in 50 mM Tris-HCl (pH 7.5) with 0.4 mg/mL RNaseA (Sigma-Aldrich), and incubated at 37°C for 1 hour. Protein digestion was carried out with 5 µL of Proteinase K (20 mg/mL, GRiSP) for 2 hours at 55°C. After centrifugation and washing with 50 mM Tris-HCl (pH 7.5), cells were diluted 10-fold in 50 mM Tris-HCl (pH 7.8) containing 1 mM SYTOX Green (Thermo Fisher) and analyzed using a Fortessa flow cytometer (BD Bioscience). DNA content per cell was quantified via the FITC channel, acquiring 10,000 events per sample. Cell cycle profiles were analyzed and visualized in Python v3.9 using the FlowCytometryTools package (https://eyurtsev.github.io/FlowCytometryTools/). The time spent in G1 and G2 phases for each genotype was estimated by deriving the the relative cell distribution across cell cycle phases from FACS data, and multiplying them by the doubling time estimated from growth curves under each glucose condition. Mean values were calculated from three independent biological replicates. Statistical analysis was performed using ANOVA and Tukey’s HSD for pairwise comparisons. A significance threshold of adjusted p-value < 0.05 was applied. All experiments were performed at the Flow Cytometry Facility of Gulbenkian Institute for Molecular Medicine.

### Cell size

Three independent replicates of ancestral WT and *ctf4Δ* strains were grown to saturation in YPD with varying glucose concentrations (0.25%, 0.5%, 2%, or 8%). Cell volume was measured using a Coulter Counter (Multisizer 4e, Beckman), which estimates particle sizes based on the Coulter principle: as a cell passes through an electrolyte-filled orifice, the impedance change is proportional to the cell volume. For measurements, 10 µL of cell culture was transferred into a cuvette containing 10 mL of conducting fluid (Isoton II solution), using a 50 µm aperture. A one-way analysis of variance (ANOVA) was used to compare cell volumes across different glucose concentrations within each genotype (WT and *ctf4Δ*). Statistical analysis was performed using ANOVA and Tukey’s HSD for pairwise comparisons. A significance threshold of adjusted p-value < 0.05 was applied.

### Fitness Assay

To assess relative fitness, strains were competed against a fluorescently labeled reference strain. The reference strain was generated by integrating the pFA6a-prACT1-yCerulean-HphMX4 plasmid, digested with AgeI, into the *ACT1* locus of the ancestral WT, enabling strong expression of the yCerulean fluorescent protein under the *ACT1* promoter. For ancestral strains, WT, *ctf4Δ*, and the reference strain were inoculated from frozen stocks into 5 mL YPD tubes and grown to saturation at 30°C. For evolved strains, 96-well plates containing frozen populations were thawed, and 10 µL of each population was inoculated into deep 96-well plates containing 1 mL of culture medium with the appropriate glucose concentration/composition. Cells were grown for 24–48 hours, depending on the passage. For the competition assay, test and reference strains were mixed in a 1 mL culture (in 96-well plates) at a ratio reflecting the expected fitness difference (e.g., 1:1 for near-equal fitness) and allowed to proliferate at 30°C for 24 hours. An initial sample (10 µL) was taken after 5 hours (day 0) and strain ratios were immediately measured using a flow cytometer (HTS, Fortessa, BD Bioscience). Cultures were then propagated with 1:1000 dilutions into fresh medium every 24 hours for other 2 days. Strain ratios and the number of generations between samples were measured at each passage.

Ratios *r* were calculated based on the number of fluorescent and non-fluorescent events detected by the flow cytometer:

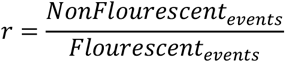

Generations between time points *g* were calculated based on total events measured at time 0 and time 24 hr:

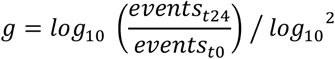

Linear regression was performed between the (*g, log*_*e*_ *r*) points relative to every sample. Relative fitness was calculated as the slope of the resulting line. The mean relative fitness was calculated from measurements obtained from at least three independent biological replicates. Relative fitness of the ancestral WT strain was used to normalize fitness across conditions. For reconstruction of adaptive mutations, the fitness effect of each reconstructed mutation in WT background was used to normalize fitness in *ctf4*Δ background. All measurements of delta (Δ) fitness represent the difference between the tested strain and *ctf4*Δ ancestor. Statistical analysis was performed using ANOVA and Tukey’s HSD for pairwise comparisons, except when mentioned otherwise. A significance threshold of adjusted p-value < 0.05 was applied. The evolved lines fitness trajectories were fitted with both a power law and linear regression and R^2^ was calculated. The power law formula was adapted from (Wiser et al., 2013) is expressed as:

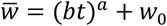

Where 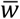 represents mean evolved fitness, *t* represents generations, *w*_0_ denotes ancestral fitness. Two parameters, *b* and *a*, were estimated based on the fit, with *b* serving as a proxy for the adaptation rate. Fit and estimation of b parameter was not possible for populations 1, 5 and 10 of evolved WT in 2% glucose. Pair-wise comparisons using Mann-Whitney with Bonferroni correction were preformed to compare estimated adaptation rate between glucose conditions. Statistical analysis to determine differences between low and high glucose in reconstructed strains was preformed using one-way ANOVA F-statistic with Bonferroni correction. All fitness experiments were performed at the Flow Cytometry Facility of Gulbenkian Institute for Molecular Medicine.

### Experimental Evolution

The 48 populations used for experimental evolution were derived from the same isogenic clone, initially inoculated in 5 mL of YPD. Evolution was performed in a high-throughput system using deep 96-well plates, each well containing 1.5 mL of medium. Ten microliters of saturated isogenic culture were inoculated into each well. Twelve parallel populations of each genotype evolved under different glucose concentrations, following a fixed layout. Plates were incubated at 30°C with 70% humidity using the Multi-Purpose Tube Rotator 5 - 80 rpm (Fisher Scientific). Plates were fixed to a modified 64-place drum tube carousel (Fisher Scientific) with a custom-made 3D printed plate holder, and rotated at 20 rpm set to the second degree of inclination available. Plates were sealed with sterile breathable rayon film (Adhesive Fil for Culture, VWR). Daily passages were conducted by diluting a defined number of cells, adjusted to maintain Ne within the same order of magnitude. Ne was estimated using the formula:

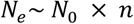

Where *N*_0_ is the initial population size calculated based on the bottle neck and final population size and *n* is the number of generations which can be calculated based using by calculating the log2 of the dilution factor (Lenski et al., 1991). A 12-channel pipette (10 µL) was used to perform dilutions, following a vertical order to avoid cross-contamination between parallel populations. The standard dilution for WT at 2% glucose was 1:1000, allowing approximately 10 generations per passage. Cell concentration and dilution rates were adjusted at passages 5, 15, 30, 60, 90, and 100. All populations were evolved for a total of 1000 generations. To prevent WT cross-contamination in *ctf4Δ* populations, G418 (200 µg/mL) was added to the medium every five passages. At every fifth passage, 100 µL of each evolving population was mixed with 100 µL of 30% (v/v) glycerol and stored at ™80°C in shallow 96-well plates sealed with Adhesive Foil (VWR), following the same layout as the evolution plates for future analysis. Due to contamination, population 8 of WT evolved in 0.5% glucose was excluded from all the analysis.

### Whole genome sequencing

Genomic DNA library preparation was performed as in (Koschwanez et al., 2013) with an Illumina Nextera DNA Library Prep Kit. Libraries were then pooled and sequenced with an Illumina NovaSeq (150 bp paired end reads). The SAMtools software (samtools.sourceforge.net) was then used to sort and index the mapped reads into a BAM file. GATK (www.broadinstitute.org/gatk) (McKenna et al., 2010) was used to realign local indels, and VarScan (varscan.sourceforge.net) was used to call variants. Mutations were found using a custom pipeline written in Python v3.9. The pipeline (github.com/koschwanez/mutantanalysis) compares variants between the reference strain, the ancestor strain, and the evolved strains. A variant that occurs between the ancestor and an evolved strain is labeled as a mutation if it either (1) causes a non-synonymous substitution in a coding sequence or (2) occurs in a regulatory region, defined as the 500 bp upstream and downstream of the coding sequence.

### Copy number variations

Whole genome sequencing and read mapping was done as previously described. The read-depths for every unique 100-bp region in the genome were then obtained by using the VarScan copy number tool. A custom pipeline written in Python v3.9 was used to visualize the genome-wide CNVs. First, the read-depths of individual 100 bp windows were normalized to the genome-wide median read-depth to control for differences in sequencing depths between samples. The coverage of the ancestor strains was then subtracted from the one of the evolved lines to reduce the noise in read depth visualization due to the repeated sequences across the genome. The resulting CNVs were smoothed across five 100 bp windows for a simpler visualization. Final CNVs were then plotted relative to their genomic coordinate at the center of the smoothed window. The custom pipeline used for the data analysis is available on GitHub (https://github.com/marcofumasoni/Fumasoni_and_Murray_2019). The total amount of segmental amplifications across the 48 populations was determined. To determine whether the 11 segmental amplifications within the region of *SCC2* was statistically significant, a binomial test was performed.

### Convergent evolution

This method was preformed based on the analysis of (Fumasoni & Murray, 2020). Briefly, it relies on the assumption that those genes that have been mutated significantly more than expected by chance alone, represent cases of convergent evolution among independent lines. The mutations affecting those genes are therefore considered putatively adaptive. The same procedure was used independently on the mutations found in WT and *ctf4*Δ evolved lines:

We first calculated per-base mutation rates as the total number of mutations in coding regions occurring in a given background (*ctf4*Δ evolved or WT evolved), divided by the size of the coding yeast genome in bp (including 1000 bp per ORF to account for regulatory regions)

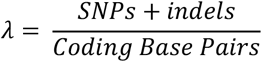

If the mutations were distributed randomly in the genome at a rate λ, the probability of finding n mutations in a given gene of length N is given by the Poisson distribution:

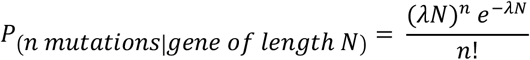

For each gene of length N, we then calculated the probability of finding n mutations if these were occurring randomly.

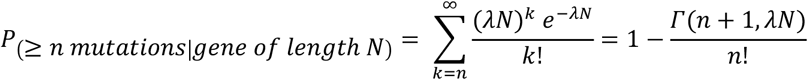

(Where *Γ* is the upper incomplete gamma function) which gives us the p-value for the comparison of the observed mutations with the null, Poisson model. In order to decrease the number of false positives, we then performed multiple-comparison corrections. Benjamini-Hochberg correction (α =0.05) was used for both the WT and *ctf4*Δ mutation dataset. Supplementary File 3 lists the mutations detected in evolved *ctf4*Δ clones, after filtering out those that occurred in genes that were significantly mutated in the WT populations. Genes significantly selected in these clones are shown in dark gray (after Benjamini-Hochberg correction with α =0.05). The custom pipeline used for the data analysis is available on GitHub.

### Mutational profile analysis

Mutational analysis was conducted using both the dataset of mutations in coding regions of *ctf4Δ* and the output from convergent evolution analyses. Mutation counts per population and genotype were analyzed to assess mutational profiles. Total and synonymous mutations were calculated separately for each genotype and glucose concentration. A Mann-Whitney U test was applied to compare mutation counts between genotypes for both total and synonymous mutations. Mutation counts were also grouped by glucose concentration for each genotype, and the Kruskal-Wallis test was test the effect of glucose in mutation counts. To assess clonality in evolved populations, read fraction was used as a metric. Read or mutation fraction represents the percentage of sequencing reads containing a specific mutation, with 100% indicating the mutation is present in all sequenced genomes. The median read fraction for each population was calculated and grouped by genotype and glucose concentration. Mann-Whitney U tests were performed to compare median read fractions between WT and *ctf4Δ*, and ANOVA with Turkey’s HSD was used to assess differences across glucose concentrations. Finally, the Kolmogorov-Smirnov test was employed to compare the distribution of mutation fractions across glucose concentrations between genotypes.

### GO enrichment analysis

To identify functions under selection that may be influenced by mutations across multiple genes, we performed GO enrichment analysis on mutations found to be positively selected in *ctf4Δ* evolved clones, using two complementary methods. First, we identified putatively adaptive GO terms using the same approach described above, but considering the GO terms, instead of genes, as the unit of our analysis. (see Supplementary File 3) Second, we analyzed the full list of mutated CDS genes in *ctf4Δ* evolved clones by constructing an interaction network in Cytoscape, using the STRING database (https://string-db.org) plug-in. This approach allowed us to identify genes that, while not individually flagged for adaptive mutations, may belong to functional modules under selection. Mutations in these genes could contribute to the overall phenotype. The interaction network was curated and visualized in Cytoscape. GO enrichment for the RNA-seq data was preformed using the GO Term Finder tool for biological process of the Saccharomyces Genome Database (https://www.yeastgenome.org/).

### RNA-seq Analysis

The RNA-seq dataset from (Warfield et al., 2022), available at the Gene Expression Omnibus under accession GEO: GSE190778, the signal per gene across samples was normalized using *Schizosaccharomyces pombe* reads mapped as spike-ins. Data counts were rounded and imported into the R programming language (v.3.6.3) for differential gene expression analysis and visualization using the DESeq2 package (v.1.31.7) (Love et al., 2014). We analyzed gene expression from three independent replicates of degron-mediated TID disruption (TID_3IAA) and corresponding controls (TID_DMSO), resulting in 4,338 genes for downstream analysis. The DESeq2 pipeline was implemented using the DESeq function, which estimates size factors with estimateSizeFactors, dispersion with estimateDispersions, and performs binomial GLM fitting and Wald statistics using nbinomWaldTest. Pairwise comparisons between TID_3IAA and TID_DMSO were tested through contrasts using the results function. Differentially expressed genes were defined by an adjusted p-value < 0.01 and an absolute log2 fold change > 1 (equivalent to a 2-fold change). Normalized gene expression counts were obtained with the counts function (using normalized = TRUE). Results from analysis are available in Supplementary File 4.

### Mediator Complex Structure Visualization and Amino Acid Substitution Modeling

Molecular graphics and analysis performed with UCSF ChimeraX, developed by the Resource for Biocomputing, Visualization, and Informatics at the University of California, San Francisco, with support from National Institutes of Health R01-GM129325 and the Office of Cyber Infrastructure and Computational Biology, National Institute of Allergy and Infectious Diseases. (Meng et al., 2023). The publicly available structure (PDB: 7UIO) was uploaded to ChimeraX for visualization and color coding. The Rotamers Structure Editing tool was used to model amino acid substitutions, using the Dunbrack rotamer library (Shapovalov & Dunbrack, 2011) was used to predict steric clashes with neighboring residues. We selected the rotamer with the highest prevalence (0.813) and minimal steric clashes to reduce the likelihood of introducing structural artifacts into the model.

## Acknowledgments

We thank Jorge Carneiro, Thomas LaBar, Isabel Gordo, Ana Garoña and Manuel Vasquez for critical reading of the manuscript; Adolfo Alsina for assistance in cell cycle data analysis; Victor Mello for assistance in protein structure analysis; All the members of the Genome Maintenance and Evolution lab for helpful discussions; Tiago Paixão and Hugo Lainé from the Advanced Data Analysis (ADA) facility of the Gulbenkian Institute for Molecular Medicine for their availability and help with RNA-seq data processing and statistical analysis; the Flow Cytometry and Genomics Facility of Gulbenkian Institute for Molecular Medicine for their technical support. M.N. acknowledges the support of a Fundação para a Ciência e a Tecnologia (FCT) doctoral fellowship (UI/BD/152252/2021). M.F. acknowledges the support of the Horizon 2020 Marie Skłodowska-Curie Actions (101030203-M.F.) and an FCT fellowship (2023.09068.CEECIND). Work in the Genome Maintained and Evolution lab was supported by FCT (2022.07846.PTDC), EMBO (5349-2023), HFSP (RGEC28/2023), the Gulbenkian Foundation (FCG) and the Gulbenkian Institute for Molecular Medicine Foundation (GIMM).

## References

Abe, T., Kawasumi, R., Giannattasio, M., Dusi, S., Yoshimoto, Y., Miyata, K., Umemura, K., Hirota, K., & Branzei, D. (2018). AND-1 fork protection function prevents fork resection and is essential for proliferation. Nature Communications, 9(1), 3091.

Alberghina, L., Smeraldi, C., Ranzi, B. M., & Porro, D. (1998). Control by nutrients of growth and cell cycle progression in budding yeast, analyzed by double-tag flow cytometry. Journal of Bacteriology, 180(15), 3864–3872.

Ardell, S. M., Martsul, A., Johnson, M. S., & Kryazhimskiy, S. (2024). Environment-independent distribution of mutational effects emerges from microscopic epistasis. Science, 386(6717), 87–92.

Beck, C., & von Meyenburg, H. K. (1968). Enzyme pattern and aerobic growth of Saccharomyces cerevisiae under various degrees of glucose limitation. Journal of Bacteriology, 96(2), 479–486.

Boer, V. M., Amini, S., & Botstein, D. (2008). Influence of genotype and nutrition on survival and metabolism of starving yeast. Proceedings of the National Academy of Sciences of the United States of America, 105(19), 6930–6935.

Bourbon, H.-M. (2008). Comparative genomics supports a deep evolutionary origin for the large, four-module transcriptional mediator complex. Nucleic Acids Research, 36(12), 3993–4008.

Boyer, S., Hérissant, L., & Sherlock, G. (2021). Adaptation is influenced by the complexity of environmental change during evolution in a dynamic environment. PLoS Genetics, 17(1), e1009314.

Brauer, M. J., Huttenhower, C., Airoldi, E. M., Rosenstein, R., Matese, J. C., Gresham, D., Boer, V. M., Troyanskaya, O. G., & Botstein, D. (2008). Coordination of growth rate, cell cycle, stress response, and metabolic activity in yeast. Molecular Biology of the Cell, 19(1), 352–367.

Carter, B. L., & Jagadish, M. N. (1978). Control of cell division in the yeast Saccharomyces cerevisiae cultured at different growth rates. Experimental Cell Research, 112(2), 373–383.

Corcoles-Saez, I., Ferat, J.-L., Costanzo, M., Boone, C. M., & Cha, R. S. (2019). Functional link between mitochondria and Rnr3, the minor catalytic subunit of yeast ribonucleotide reductase. Microbial Cell (Graz, Austria), 6(6), 286–294.

Costanzo, M., Hou, J., Messier, V., Nelson, J., Rahman, M., VanderSluis, B., Wang, W., Pons, C., Ross, C., Ušaj, M., San Luis, B.-J., Shuteriqi, E., Koch, E. N., Aloy, P., Myers, C. L., Boone, C., & Andrews, B. (2021). Environmental robustness of the global yeast genetic interaction network. Science, 372(6542), eabf8424–eabf8424.

Couce, A., & Tenaillon, O. A. (2015). The rule of declining adaptability in microbial evolution experiments. Frontiers in Genetics, 6(MAR), 99.

Crow, J. F., & Kimura, M. (1970). An introduction to population genetics theory. Blackburn Press.

Debray, R., De Luna, N., & Koskella, B. (2022). Historical contingency drives compensatory evolution and rare reversal of phage resistance. Molecular Biology and Evolution, 39(9), msac182.

Desai, M. M., Fisher, D. S., & Murray, A. W. (2007). The speed of evolution and maintenance of variation in asexual populations. Current Biology: CB, 17(5), 385–394.

Diaz-Colunga, J., Skwara, A., Gowda, K., Diaz-Uriarte, R., Tikhonov, M., Bajic, D., & Sanchez, A. (2023). Global epistasis on fitness landscapes. Philosophical Transactions of the Royal Society of London. Series B, Biological Sciences, 378(1877). 10.1098/rstb.2022.0053

Fagny, M., & Austerlitz, F. (2021). Polygenic adaptation: Integrating population genetics and gene regulatory networks. Trends in Genetics: TIG, 37(7), 631–638.

Ferrari, E., Bruhn, C., Peretti, M., Cassani, C., Carotenuto, W. V., Elgendy, M., Shubassi, G., Lucca, C., Bermejo, R., Varasi, M., Minucci, S., Longhese, M. P., & Foiani, M. (2017). PP2A Controls Genome Integrity by Integrating Nutrient-Sensing and Metabolic Pathways with the DNA Damage Response. Molecular Cell, 67(2), 266-281.e4.

Filteau, M., Hamel, V., Pouliot, M., Gagnon-Arsenault, I., Dubé, A. K., & Landry, C. R. (2015). Evolutionary rescue by compensatory mutations is constrained by genomic and environmental backgrounds. Molecular Systems Biology, 11(10), 832–832.

Fumasoni, M. (2020). Tell Me Where You’ve Been and I’ll Tell You How You’ll Evolve. MBio, 11(5), 2043–2063.

Fumasoni, M., & Murray, A. W. (2020). The evolutionary plasticity of chromosome metabolism allows adaptation to constitutive DNA replication stress. ELife, 9, 1–28.

Fumasoni, M., & Murray, A. W. (2021). Ploidy and recombination proficiency shape the evolutionary adaptation to constitutive DNA replication stress. PLoS Genetics, 17, e1009875.

Fumasoni, M., Zwicky, K., Vanoli, F., Lopes, M., & Branzei, D. (2015). Error-Free DNA Damage Tolerance and Sister Chromatid Proximity during DNA Replication Rely on the Polα/Primase/Ctf4 Complex. Molecular Cell, 57(5), 812– 823.

Gaillard, H., García-Muse, T., & Aguilera, A. (2015). Replication stress and cancer. Nature Reviews. Cancer, 15(5), 276–280.

Gambus, A., Van Deursen, F., Polychronopoulos, D., Foltman, M., Jones, R. C., Edmondson, R. D., Calzada, A., & Labib, K. (2009). A key role for Ctf4 in coupling the MCM2-7 helicase to DNA polymerase α within the eukaryotic replisome. The EMBO Journal, 28(19), 2992–3004.

Gorbea Colón, J. J., Palao, L., 3rd, Chen, S.-F., Kim, H. J., Snyder, L., Chang, Y.-W., Tsai, K.-L., & Murakami, K. (2023). Structural basis of a transcription pre-initiation complex on a divergent promoter. Molecular Cell, 83(4), 574-588.e11.

Gorter, F. A., Derks, M. F. L., van den Heuvel, J., Aarts, M. G. M., Zwaan, B. J., de Ridder, D., & de Visser, J. A. G. M. (2017). Genomics of adaptation depends on the rate of environmental change in experimental yeast populations. Molecular Biology and Evolution. 10.1093/molbev/msx185

He, Z., Houghton, P. J., Williams, T. M., & Shen, C. (2021). Regulation of DNA duplication by the mTOR signaling pathway. Cell Cycle (Georgetown, Tex.), 20(8), 742–751.

Hietpas, R. T., Bank, C., Jensen, J. D., & Bolon, D. N. A. (2013). Shifting fitness landscapes in response to altered environments: Fitness landscapes in altered environments. Evolution; International Journal of Organic Evolution, 67(12), 3512–3522.

Hsieh, Y. Y. P., Makrantoni, V., Robertson, D., Marston, A. L., & Murray, A. W. (2020). Evolutionary repair: Changes in multiple functional modules allow meiotic cohesin to support mitosis. PLoS Biology, 18(3), 1–35.

Johnson, M. S., Reddy, G., & Desai, M. M. (2023). Epistasis and evolution: recent advances and an outlook for prediction. BMC Biology, 21(1), 120.

Johnston, G. C., Ehrhardt, C. W., Lorincz, A., & Carter, B. L. (1979). Regulation of cell size in the yeast Saccharomyces cerevisiae. Journal of Bacteriology, 137(1), 1–5.

Johnston, G. C., Pringle, J. R., & Hartwell, L. H. (1977). Coordination of growth with cell division in the yeast Saccharomyces cerevisiae. Experimental Cell Research, 105(1), 79–98.

Johnston, G. C., Singer, R. A., Sharrow, S. O., & Slater, M. L. (1980). Cell division in the yeast Saccharomyces cerevisiae growing at different rates. Microbiology, 118(2), 479–484.

Kim, Y.-J., Björklund, S., Li, Y., Sayre, M. H., & Kornberg, R. D. (1994). A multiprotein mediator of transcriptional activation and its interaction with the C-terminal repeat domain of RNA polymerase II. Cell, 77(4), 599–608.

Kimura, M. (1985). The role of compensatory neutral mutations in molecular evolution. Journal of Genetics, 64(1), 7–19.

Koschwanez, J. H., Foster, K. R., & Murray, A. W. (2013). Improved use of a public good selects for the evolution of undifferentiated multicellularity. ELife, 2013(2), e00367.

Kryazhimskiy, S., Rice, D. P., Jerison, E. R., & Desai, M. M. (2014). Microbial evolution. Global epistasis makes adaptation predictable despite sequence-level stochasticity. Science, 344(6191), 1519–1522.

Kwan, E. X., Foss, E. J., Tsuchiyama, S., Alvino, G. M., Kruglyak, L., Kaeberlein, M., Raghuraman, M. K., Brewer, B. J., Kennedy, B. K., Bedalov, A., Kaeberlein, M., Mortimer, R. K., Johnston, J. R., Steinkraus, K. A., Kaeberlein, M., Kennedy, B. K., Kennedy, B. K., Gotta, M., Sinclair, D. A., … Fangman, W. L. (2013). A Natural Polymorphism in rDNA Replication Origins Links Origin Activation with Calorie Restriction and Lifespan. PLoS Genetics, 9(3), e1003329.

Laan, L., Koschwanez, J. H., & Murray, A. W. (2015). Evolutionary adaptation after crippling cell polarization follows reproducible trajectories. ELife, 4(4:e09638), e09638.

LaBar, T., Phoebe Hsieh, Y.-Y., Fumasoni, M., & Murray, A. W. (2020). Evolutionary Repair Experiments as a Window to the Molecular Diversity of Life. Current Biology: CB, 30(10), R565–R574.

Lamm, N., Rogers, S., & Cesare, A. J. (2019). The mTOR pathway: Implications for DNA replication. Progress in Biophysics and Molecular Biology, 147, 17–25.

Lässig, M., Mustonen, V., & Walczak, A. M. (2017). Predicting evolution. Nature Ecology & Evolution 2017 1:3, 1(3), 1–9.

Lenski, R. E., Rose, M. R., Simpson, S. C., & Tadler, S. C. (1991). Long-Term Experimental Evolution in Escherichia coli. I. Adaptation and Divergence During. The American Naturalist, 138(November), 1315–1341.

Love, M. I., Huber, W., & Anders, S. (2014). Moderated estimation of fold change and dispersion for RNA-seq data with DESeq2. Genome Biology, 15(12), 550.

Lynch, M., Conery, J., & Burger, R. (1995). Mutation Accumulation and the Extinction of Small Populations. The American Naturalist, 146(4), 489–518.

Macheret, M., & Halazonetis, T. D. (2015). DNA replication stress as a hallmark of cancer. Annual Review of Pathology: Mechanisms of Disease, 10, 425–448.

McCutchan, T. F., Rathore, D., & Li, J. (2004). Compensatory evolution in the human malaria parasite Plasmodium ovale. Genetics, 166(1), 637–640.

McKenna, A., Hanna, M., Banks, E., Sivachenko, A., Cibulskis, K., Kernytsky, A., Garimella, K., Altshuler, D., Gabriel, S., Daly, M., & DePristo, M. A. (2010). The Genome Analysis Toolkit: a MapReduce framework for analyzing next-generation DNA sequencing data. Genome Research, 20(9), 1297–1303.

Meng, E. C., Goddard, T. D., Pettersen, E. F., Couch, G. S., Pearson, Z. J., Morris, J. H., & Ferrin, T. E. (2023). UCSF ChimeraX: Tools for structure building and analysis. Protein Science: A Publication of the Protein Society, 32(11), e4792.

Miles, J., & Formosa, T. (1992). Protein affinity chromatography with purified yeast DNA polymerase alpha detects proteins that bind to DNA polymerase. Proceedings of the National Academy of Sciences of the United States of America, 89(4), 1276–1280.

Myers, L. C., Gustafsson, C. M., Bushnell, D. A., Lui, M., Erdjument-Bromage, H., Tempst, P., & Kornberg, R. D. (1998). The Med proteins of yeast and their function through the RNA polymerase II carboxy-terminal domain. Genes & Development, 12(1), 45–54.

Natalino, M., & Fumasoni, M. (2023). Experimental approaches to study evolutionary cell biology using yeasts. Yeast. 10.1002/yea.3848

Papp, B., Notebaart, R. A., & Pál, C. (2011). Systems-biology approaches for predicting genomic evolution. Nature Reviews. Genetics, 12(9), 591–602.

Pavani, M., Bonaiuti, P., Chiroli, E., Gross, F., Natali, F., Macaluso, F., Póti, Á., Pasqualato, S., Farkas, Z., Pompei, S., Cosentino Lagomarsino, M., Rancati, G., Szüts, D., & Ciliberto, A. (2021). Epistasis, aneuploidy, and functional mutations underlie evolution of resistance to induced microtubule depolymerization. The EMBO Journal, 40(22), e108225.

Persi, E., Wolf, Y. I., Horn, D., Ruppin, E., Demichelis, F., Gatenby, R. A., Gillies, R. J., & Koonin, E. V. (2021). Mutation-selection balance and compensatory mechanisms in tumour evolution. Nature Reviews. Genetics, 22(4), 251–262.

Poli, J., Tsaponina, O., Crabbé, L., Keszthelyi, A., Pantesco, V., Chabes, A., Lengronne, A., Pasero, P., Alabert, C., Bianco, J. N., Pasero, P., Alcasabas, A. A., Osborn, A. J., Bachant, J., Hu, F., Werler, P. J., Bousset, K., Furuya, K., Diffley, J. F., … Rothstein, R. (2012). dNTP pools determine fork progression and origin usage under replication stress. The EMBO Journal, 31(4), 883–894.

Reddy, G., & Desai, M. M. (2021). Global epistasis emerges from a generic model of a complex trait. ELife, 10, e64740.

Sakai, A., Shimizu, Y., & Hishinuma, F. (1988). Isolation and characterization of mutants which show an oversecretion phenotype in Saccharomyces cerevisiae. Genetics, 119(3), 499–506.

Salim, D., Bradford, W. D., Freeland, A., Cady, G., Wang, J., Pruitt, S. C., & Gerton, J. L. (2017). DNA replication stress restricts ribosomal DNA copy number. PLoS Genetics, 13(9), e1007006.

Saltz, J. B., Bell, A. M., Flint, J., Gomulkiewicz, R., Hughes, K. A., & Keagy, J. (2018). Why does the magnitude of genotype-by-environment interaction vary? Ecology and Evolution, 8(12), 6342–6353.

Samora, C. P., Saksouk, J., Goswami, P., Wade, B. O., Singleton, M. R., Bates, P. A., Lengronne, A., Costa, A., Uhlmann, F., Borges, V., Smith, D. J., Whitehouse, I., Uhlmann, F., Calì, F., Bharti, S. K., Perna, R. D., Brosh, R. M., Pisani, F. M., Chan, K.-L., … Masai, H. (2016). Ctf4 Links DNA Replication with Sister Chromatid Cohesion Establishment by Recruiting the Chl1 Helicase to the Replisome. Molecular Cell, 63(3), 371–384.

Schenk, M. F., Zwart, M. P., Hwang, S., Ruelens, P., Severing, E., Krug, J., & de Visser, J. A. G. M. (2022). Population size mediates the contribution of high-rate and large-benefit mutations to parallel evolution. Nat Ecol Evol. https://www.biorxiv.org/content/10.1101/2021.02.02.429281v1?s=09

Schonbrun, M., Kolesnikov, M., Kupiec, M., & Weisman, R. (2013). TORC2 is required to maintain genome stability during S phase in fission yeast. The Journal of Biological Chemistry, 288(27), 19649–19660.

Shapovalov, M. V., & Dunbrack, R.L., Jr. (2011). A smoothed backbone-dependent rotamer library for proteins derived from adaptive kernel density estimates and regressions. Structure (London, England: 1993), 19(6), 844–858.

Shen, C., Lancaster, C. S., Shi, B., Guo, H., Thimmaiah, P., & Bjornsti, M.-A. (2007). TOR Signaling Is a Determinant of Cell Survival in Response to DNA Damage. Molecular and Cellular Biology, 27(20), 7007–7017.

Shimada, K., Filipuzzi, I., Stahl, M., Helliwell, S. B., Studer, C., Hoepfner, D., Seeber, A., Loewith, R., Movva, N. R., & Gasser, S. M. (2013). TORC2 Signaling Pathway Guarantees Genome Stability in the Face of DNA Strand Breaks. Molecular Cell, 51(6), 829–839.

Silander, O. K., Tenaillon, O., & Chao, L. (2007). Understanding the evolutionary fate of finite populations: the dynamics of mutational effects. PLoS Biology, 5(4), e94.

Silvera, D., Ernlund, A., Arju, R., Connolly, E., Volta, V., Wang, J., & Schneider, R. J. (2017). mTORC1 and -2 Coordinate Transcriptional and Translational Reprogramming in Resistance to DNA Damage and Replicative Stress in Breast Cancer Cells. Molecular and Cellular Biology, 37(5).

Slater, M. L., Sharrow, S. O., & Gart, J. J. (1977). Cell cycle of Saccharomycescerevisiae in populations growing at different rates. Proceedings of the National Academy of Sciences of the United States of America, 74(9), 3850–3854.

Soutourina, J. (2018). Transcription regulation by the Mediator complex. Nature Reviews. Molecular Cell Biology, 19(4), 262– 274.

Syring, I., Klümper, N., Offermann, A., Braun, M., Deng, M., Boehm, D., Queisser, A., von Mässenhausen, A., Brägelmann, J., Vogel, W., Schmidt, D., Majores, M., Schindler, A., Kristiansen, G., Müller, S. C., Ellinger, J., Adler, D., & Perner, S. (2016). Comprehensive analysis of the transcriptional profile of the Mediator complex across human cancer types. Oncotarget, 7(17), 23043–23055.

Szamecz, B., Boross, G., Kalapis, D., Kovács, K., Fekete, G., Farkas, Z., Lázár, V., Hrtyan, M., Kemmeren, P., Groot Koerkamp, M.J.A., Rutkai, E., Holstege, F. C. P., Papp, B., & Pál, C. (2014). The Genomic Landscape of Compensatory Evolution. PLoS Biology, 12(8), e1001935.

Takeuchi, Y., Horiuchi, T., & Kobayashi, T. (2003). Transcription-dependent recombination and the role of fork collision in yeast rDNA. Genes & Development, 17(12), 1497–1506.

Talavera, R. A., Prichard, B. E., Sommer, R. A., Leitao, R. M., Sarabia, C. J., Hazir, S., Paulo, J. A., Gygi, S. P., & Kellogg, D. R. (2024). Cell growth and nutrient availability control the mitotic exit signaling network in budding yeast. The Journal of Cell Biology, 223(8), e202305008.

Tanaka, H., Katou, Y., Yagura, M., Saitoh, K., Itoh, T., Araki, H., Bando, M., & Shirahige, K. (2009). Ctf4 coordinates the progression of helicase and DNA polymerase alpha. Genes to Cells: Devoted to Molecular & Cellular Mechanisms, 14(7), 807–820.

Thompson, C. M., & Young, R. A. (1995). General requirement for RNA polymerase II holoenzymes in vivo. Proceedings of the National Academy of Sciences of the United States of America, 92(10), 4587–4590.

Torrence, M. E., & Manning, B. D. (2018). Nutrient Sensing in Cancer. Annual Review of Cancer Biology, 2(1), 251–269.

Tsaponina, O., Barsoum, E., Åström, S. U., & Chabes, A. (2011). Ixr1 is required for the expression of the ribonucleotide reductase Rnr1 and maintenance of dNTP pools. PLoS Genetics, 7(5). 10.1371/journal.pgen.1002061

Turner, J. J., Ewald, J. C., & Skotheim, J. M. (2012). Cell size control in yeast. Current Biology: CB, 22(9), R350–9.

Van den Bergh, B., Swings, T., Fauvart, M., & Michiels, J. (2018). Experimental Design, Population Dynamics, and Diversity in Microbial Experimental Evolution. Microbiology and Molecular Biology Reviews: MMBR, 82(3), e00008–18.

Villa, F., Simon, A. C., Ortiz Bazan, M. A., Kilkenny, M. L., Wirthensohn, D., Wightman, M., Matak-Vinkovíc, D., Pellegrini, L., Labib, K., Araki, H., Hamatake, R. K., Johnston, L. H., Sugino, A., Borges, V., Smith, D. J., Whitehouse, I., Uhlmann, F., Buser, R., Kellner, V., … Dutta, A. (2016). Ctf4 Is a Hub in the Eukaryotic Replisome that Links Multiple CIP-Box Proteins to the CMG Helicase. Molecular Cell, 63(3), 385–396.

Warfield, L., Donczew, R., Mahendrawada, L., & Hahn, S. (2022). Yeast Mediator facilitates transcription initiation at most promoters via a Tail-independent mechanism. Molecular Cell, 82(21), 4033-4048.e7.

Weisman, R., Cohen, A., & Gasser, S. M. (2014). TORC 2—a new player in genome stability. EMBO Molecular Medicine, 6(8), 995–1002.

Wiser, M. J., Ribeck, N., & Lenski, R. E. (2013). Long-term dynamics of adaptation in asexual populations. Science, 342(6164), 1364–1367.

Wortel, M. T., Agashe, D., Bailey, S. F., Bank, C., Bisschop, K., Blankers, T., Cairns, J., Colizzi, E. S., Cusseddu, D., & Desai, M. M. (2021). Towards evolutionary predictions: current promises and challenges. http://ecoevorxiv.org/repository/view/3992/

Wright, S. (1931). Evolution in Mendelian populations. Genetics, 16(2), 97–159.

Yang, Q. E., MacLean, C., Papkou, A., Pritchard, M., Powell, L., Thomas, D., Andrey, D. O., Li, M., Spiller, B., Yang, W., & Walsh, T. R. (2020). Compensatory mutations modulate the competitiveness and dynamics of plasmid-mediated colistin resistance in Escherichia coli clones. The ISME Journal, 14(3), 861–865.

Yuan, Z., Georgescu, R., Santos, R. de L. A., Zhang, D., Bai, L., Yao, N. Y., Zhao, G., O’Donnell, M. E., & Li, H. (2019). Ctf4 organizes sister replisomes and Pol α into a replication factory. ELife, 8. 10.7554/eLife.47405

