## Supplementary Figures and Figure Legends for "Compensatory Evolution to DNA Replication Stress is Robust to Nutrient Availability"

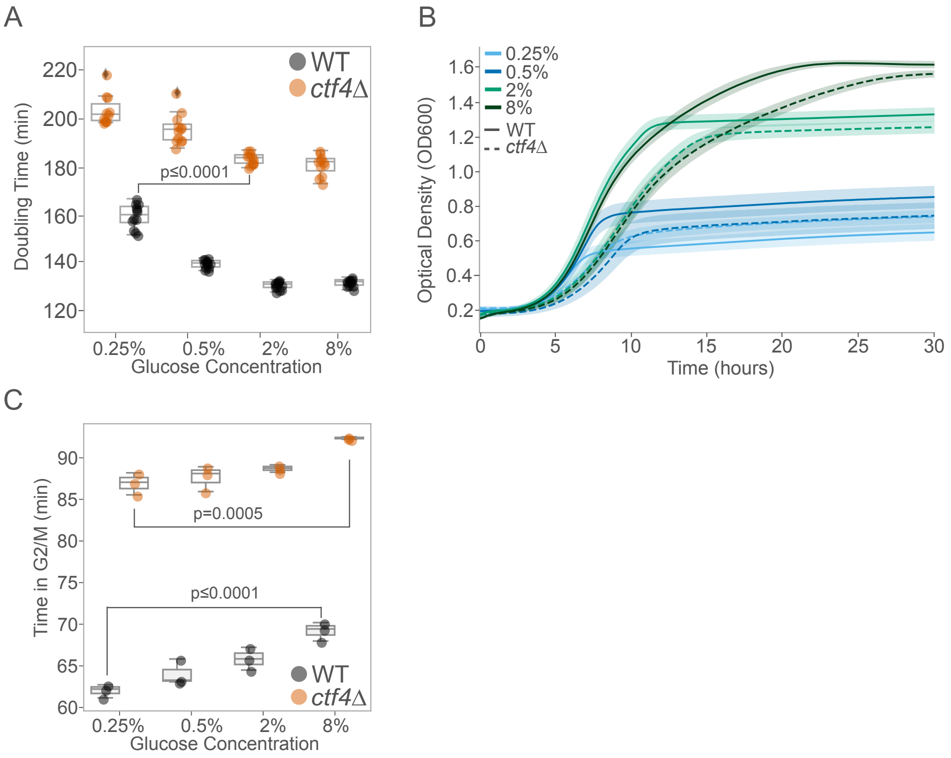


**Figure S1.** *Glucose concentration impacts growth dynamics in the presence of DNA replication stress*. (A) Population doubling time (min) of ancestral WT (black) and *ctf4*Δ mutant (orange) across different glucose concentrations. Box plots show median, IQR, and whiskers extending to 1.5×IQR, with individual data points beyond whiskers as outliers. (B) Growth curve of ancestral WT (solid line) and *ctf4*Δ (dashed line) over 30h. Colors represent different glucose concentrations: light blue (0.25%), dark blue (0.5%), light green (2%), and dark green (8%). Bold lines indicate mean growth, shaded areas represent SD. (C) Time spent (minutes) in G2/M phase for ancestral WT and *ctf4*Δ, across different glucose concentrations, estimated from DNA content and doubling times (see Materials and Methods). Detailed statistical analysis and underlying data for this figure are provided in Supplementary File 1.

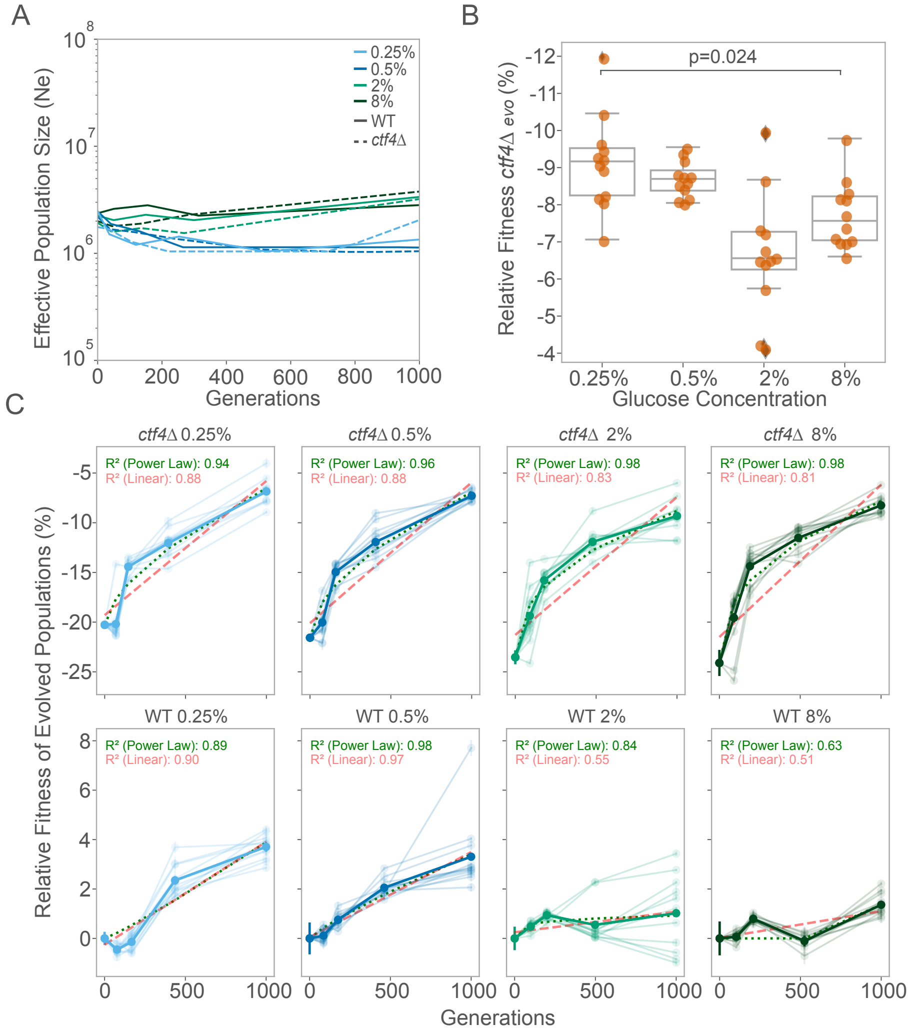


**Figure S2**. *Evolutionary dynamics under different glucose concentrations*. (A) Estimated Ne across generations. Solid and dashed lines represent WT and *ctf4*Δ adjusted Ne values across generations, respectively. (B) Fitness of *ctf4*Δ evolved populations at 1000 generations relative to WT reference. Each data point represents one parallelly evolved population. Box plots show median, IQR, and whiskers extending to 1.5×IQR, with individual data points beyond whiskers considered outliers. (C) Fitness trajectories of *ctf4*Δ (upper panel) and WT (bottom panel) populations evolved across varying glucose concentrations were fit using both a power law (green pointed line) and linear (red dashed line) regression. Individual population data (shaded lines) and mean fitness values (solid lines) were plotted over generations, with error bars representing SD. Colors represent glucose concentrations: light blue (0.25%), dark blue (0.5%), light green (2%), dark green (8%). The represented values of $R^2$ are a mean from the individual populations. The detailed statistical analysis and underlying data for this figure, as well as the parameters estimated from both linear regression and power law are available in Supplementary File 2.

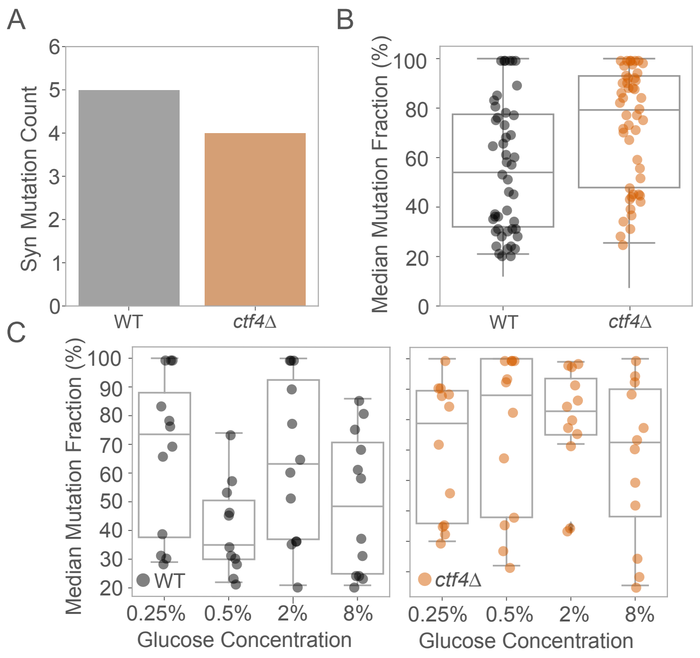


**Figure S3.** *Mutational counts*. (A) Total counts of synonymous (syn) mutations detected in evolved WT and *ctf4*Δ populations. (B) Median mutation fraction (%) in CDS for evolved WT (black) and *ctf4*Δ (orange) populations at generation 1000. Box plots show median, IQR, and whiskers extending to 1.5×IQR, with individual data points beyond whiskers considered outliers. (C) Median mutation fraction (%) per glucose concentration, per genotype (WT (left) and *ctf4*Δ (right) at generation 1000. Detailed statistical analysis and underlying data for this figure are provided in Supplementary File 3.

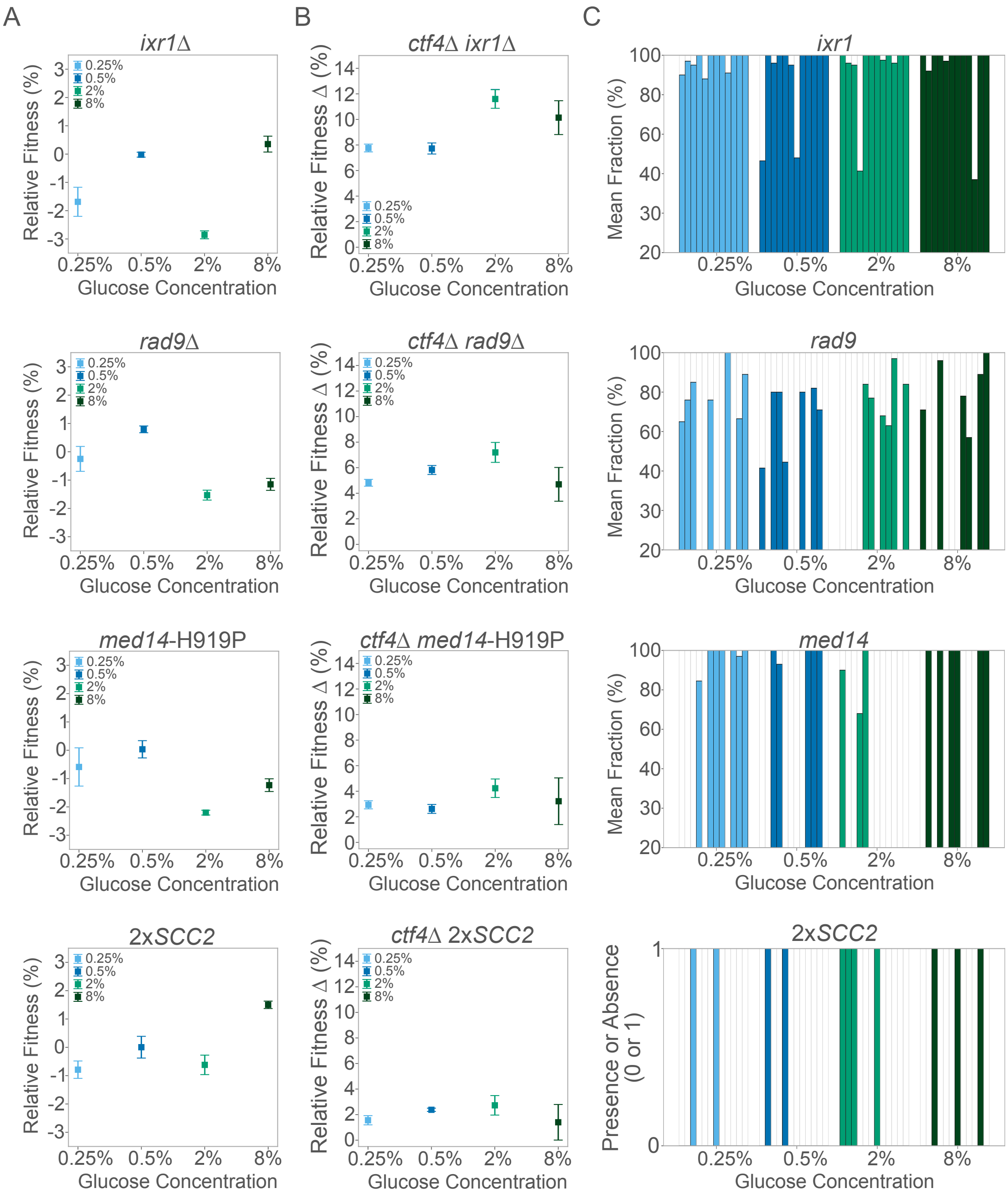


**Figure S4**. *Fitness of reconstructed strains*. (A) Mean relative fitness of reconstructed putative adaptive mutations in WT background. Error bars represent SD. Colors indicate glucose concentrations: light blue (0.25%), dark blue (0.5%), light green (2%), and dark green (8%). (B) Changes in mean relative fitness (Δ) of ancestral *ctf4*Δ clones carrying reconstructed putative adaptive mutations. Error bars represent standard deviation, with errors propagated from the two fitness measurements used to calculate delta (Δ). (C) Frequencies of adaptive mutations across glucose concentrations at 1000 generations. Each bar represents the 12 parallel populations evolved in each glucose concentration, by order (1 to 12). Alleles frequencies (mean fraction) in populations were derived from deep sequencing data of genomic DNA extracted from a population sample. . Detailed statistical analysis and underlying data for this figure are provided in Supplementary File 3 and 5.
